# Chromatin sensing by the auxiliary domains of KDM5C regulates its demethylase activity and is disrupted by X-linked intellectual disability mutations

**DOI:** 10.1101/2022.01.13.476263

**Authors:** Fatima S. Ugur, Mark J. S. Kelly, Danica Galonić Fujimori

## Abstract

The H3K4me3 chromatin modification, a hallmark of promoters of actively transcribed genes, is dynamically removed by the KDM5 family of histone demethylases. The KDM5 demethylases have a number of accessory domains, two of which, ARID and PHD1, lie between the segments of the catalytic domain. KDM5C, which has a unique role in neural development, harbors a number of mutations adjacent to its accessory domains that cause X-linked intellectual disability (XLID). The roles of these accessory domains remain unknown, limiting an understanding of how XLID mutations affect KDM5C activity. Through *in vitro* binding and kinetic studies using nucleosomes, we find that while the ARID domain is required for efficient nucleosome demethylation, the PHD1 domain alone has an inhibitory role in KDM5C catalysis. In addition, the unstructured linker region between the ARID and PHD1 domains interacts with PHD1 and is necessary for nucleosome binding. Our data suggests a model in which the PHD1 domain inhibits DNA recognition by KDM5C. This inhibitory effect is relieved by the H3 tail, enabling recognition of flanking DNA on the nucleosome. Importantly, we find that XLID mutations adjacent to the ARID and PHD1 domains break this regulation by enhancing DNA binding, resulting in the loss of specificity of substrate chromatin recognition and rendering demethylase activity lower in the presence of flanking DNA. Our findings suggest a model by which specific XLID mutations could alter chromatin recognition and enable euchromatin-specific dysregulation of demethylation by KDM5C.

## INTRODUCTION

The methylation of lysine 4 on histone H3 is a chromatin modification found on euchromatin, where H3K4 trimethylation (H3K4me3) is present at gene promoter regions associated with active transcription, and where H3K4 monomethylation (H3K4me1) is found at active enhancer regions [1]. While H3K4me1/2 is demethylated by the KDM1/LSD family, H3K4me1/2/3 is dynamically regulated by the KDM5/JARID1 subfamily of Jumonji histone demethylases [2–7]. This demethylase family harbors unique auxiliary domains in addition to its catalytic domain comprised of the JmjN and JmjC segments that form a composite active site for demethylation [8,9]. KDM5A (RBP2, JARID1A), KDM5B (PLU-1, JARID1B), KDM5C (SMCX, JARID1C), and KDM5D (SMCY, JARID1D) all contain an AT-rich interaction domain (ARID), C_5_HC_2_ zinc finger domain (ZnF), and 2-3 plant homeodomains (PHD1-3). Unique to the KDM5 family is the insertion of the ARID and PHD1 domains between the JmjN and JmjC segments of the catalytic domain, and ARID and PHD1 are required for demethylase activity *in vivo* [4,10– 13]. ARID domains are DNA binding domains, and ARID of KDM5A/B has been shown to bind to GC-rich DNA [13–15]. PHD domains are H3K4 methylation reader domains with varying specificity towards unmethylated and methylated H3K4 states [16–22]. PHD1 of KDM5A/B preferentially binds the unmethylated H3 tail, and this recognition of the demethylation product allosterically stimulates demethylase activity of KDM5A *in vitro* [23–28]. In contrast, the ARID and PHD1 domains have not been extensively studied in KDM5C, which possesses a unique function in neural development and has nonredundant demethylase activity [2,29].

KDM5C is ubiquitously expressed but has highest expression levels in the brain [30,31]. This demethylase is important for neural development and dendrite morphogenesis, and KDM5C knockout mice have abnormal dendritic branching and display memory defects, impaired social behavior, and aggression [2,29]. KDM5C fine-tunes the expression of neurodevelopmental genes, as gene expression levels only change less than 2 fold upon knockout of KDM5C in mice [29,32]. KDM5C localizes to enhancers in addition to promoter regions and has been shown to also demethylate spurious H3K4me3 at enhancers during neuronal maturation [29,32–34]. In line with its neurodevelopmental function, a number of missense and nonsense mutations that cause X-linked intellectual disability (XLID) are found throughout KDM5C [31,35–39]. As KDM5C is located on the X-chromosome and the Y paralog KDM5D cannot compensate for its function, males with KDM5C XLID mutations are primarily affected with a range of mild to severe symptoms of limitations in cognition, memory, and adaptive behavior [30,31,37,38,40]. Some functionally characterized mutations have been shown to reduce demethylase activity despite not occurring in the catalytic domains, and a select few mutations have been shown to not affect demethylase activity, disrupting nonenzymatic functions instead [2,11,39,41,42]. The consequences of these XLID mutations on KDM5C at its target regions within chromatin and their impact on gene expression during neural development is not fully understood. Interestingly, a number of XLID mutations are present throughout and in between the accessory domains of KDM5C, suggesting potential disruption of their regulatory functions. The impact of these mutations on demethylase regulation is hindered by the limited understanding of the accessory domain roles in KDM5C.

Here, we sought to determine the functions of the ARID and PHD1 auxiliary domains in KDM5C and evaluate whether these functions might be disrupted by XLID mutations. We approached these questions by interrogating the recognition and demethylation of nucleosomes by KDM5C, as nucleosome substrates enable extended interactions by multiple domains of the demethylase. Our findings reveal that the ARID and PHD1 domains, as well as the linker between them, regulate nucleosome demethylation and chromatin recognition by KDM5C. We find that DNA recognition by ARID contributes to nucleosome demethylation but not nucleosome binding, which is instead driven by the unstructured linker between ARID and PHD1. In contrast, we find that PHD1 inhibits demethylation. Furthermore, we find that XLID mutations near these regulatory domains disrupt interdomain interactions and enhance affinity towards nucleosomes, resulting in nonproductive chromatin binding and inhibition of demethylation in the presence of flanking DNA. Our findings define functional roles of the ARID and PHD1 domains in the regulation of KDM5C and provide rationale for disruption of this regulation by mutations in X-linked intellectual disability.

## RESULTS

### ARID & PHD1 region contributes to productive nucleosome demethylation

Previous work has demonstrated that KDM5C is capable of demethylating H3K4me3 peptides and that the catalytic JmjN-JmjC domain and zinc finger domain are necessary for demethylase activity [2,8,42]. To evaluate the contributions of the ARID and PHD1 domains, we sought to interrogate the recognition and demethylation of nucleosomes, given the expected interactions of these domains with DNA and histone tails, respectively. We utilized an N-terminal fragment of KDM5C containing the residues 1 to 839 necessary to monitor demethylation *in vitro* (KDM5C^1-839^), as well as an analogous construct where the ARID and PHD1 region (residues 83 to 378) is replaced by a short linker (KDM5C^1-839^ ΔAP) (Figure 1A) [8]. We measured binding affinities of these constructs to both unmodified and substrate H3K4me3 core nucleosomes containing 147 bp DNA by electrophoretic mobility shift assay. KDM5C binds nucleosomes with weak affinity and with approximately a two fold affinity gain towards substrate nucleosomes, with *K*_d_^app^ of ∼7 μM for the H3K4me3 nucleosome and ∼13 μM for the unmodified nucleosome (Figure 1B). Surprisingly, the ARID and PHD1 domains have a modest contribution to nucleosome binding, as KDM5C^1-839^ ΔAP displays only a ∼3 fold reduction in nucleosome affinity and retains the two fold preference towards the substrate nucleosome (Figure 1B). The absence of a significant enhancement of nucleosome binding through ARID and PHD1 domain-mediated interactions suggests a more complex role of these domains rather than simply facilitating chromatin recruitment.

**Figure 1.**
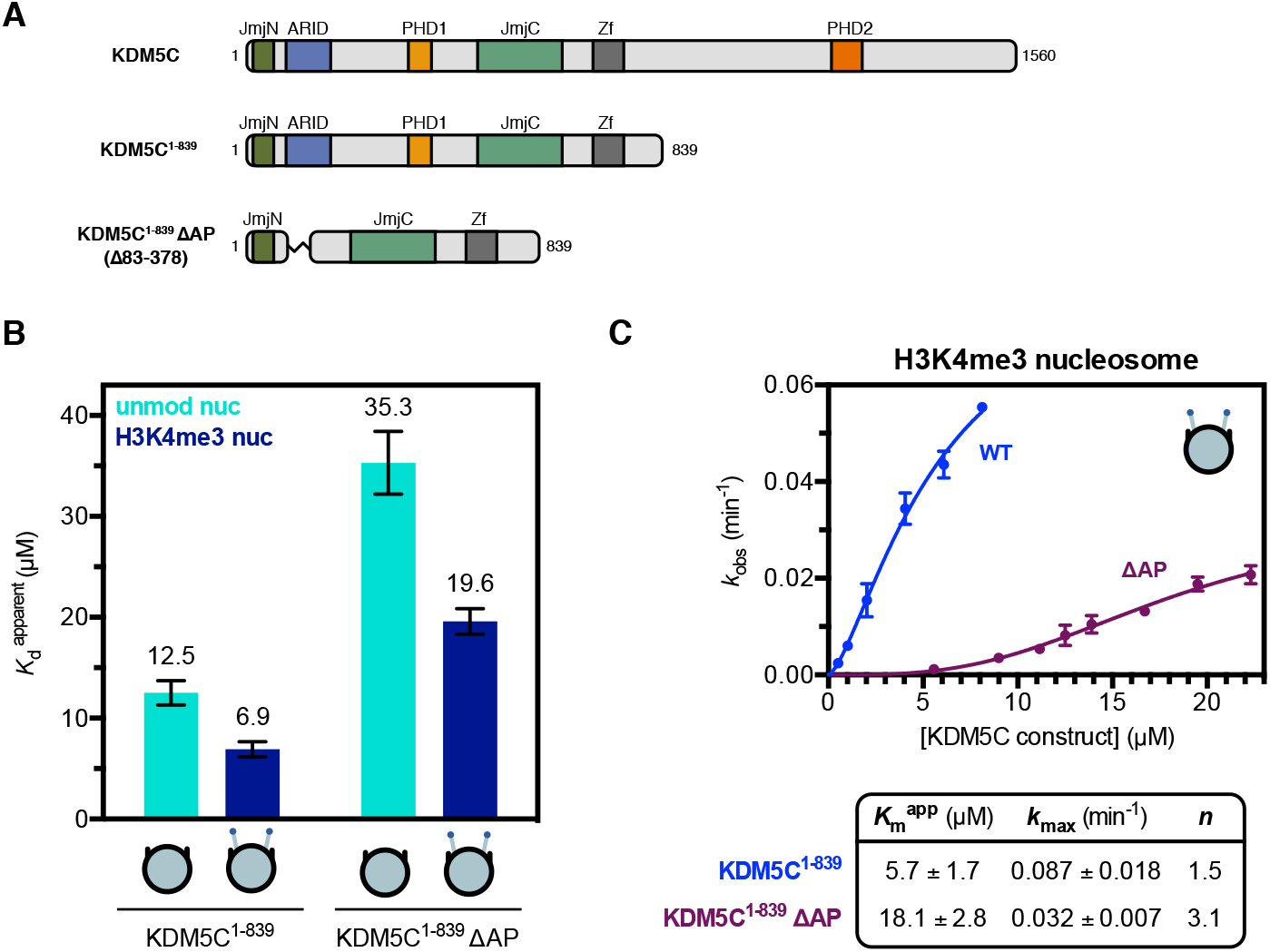
The ARID & PHD1 region of KDM5C contributes to efficient nucleosome demethylation and has a modest contribution to nucleosome binding. **(A)** Domain architecture of KDM5C and KDM5C constructs used in this study. **(B)** Unmodified and substrate nucleosome binding by KDM5C constructs with apparent dissociation constants (*K*_d_^app^) measured by EMSA (binding curves in Figure S1B). Due to unattainable saturation of binding for the unmodified nucleosome, a lower limit for the dissociation constant is presented. **(C)** Demethylation kinetics of the H3K4me3 substrate nucleosome by KDM5C constructs under single turnover conditions (enzyme in excess of substrate). Observed rates are fit to a cooperative kinetic model, with *n* denoting the Hill coefficient. Representative kinetic traces used to determine observed demethylation rates are in Figure S1C. All error bars represent SEM of at least three independent experiments (*n* ≥ 3).

We next interrogated the demethylase activity of KDM5C towards the H3K4me3 substrate nucleosome *in vitro* by utilizing a TR-FRET based kinetic assay that detects formation of the H3K4me1/2 product nucleosome. In order to measure true catalytic rates (*k*_max_), demethylation was performed under single turnover conditions with enzyme in excess [43]. We find that KDM5C^1-839^ demethylates the substrate nucleosome with an observed catalytic rate of ∼0.09 min^−1^ and KDM5C^1-839^ ΔAP with a 3-fold lower catalytic rate of ∼0.03 min^−1^ (Figure 1C), indicating that the ARID and PHD1 region contributes to productive catalysis on nucleosomes. The contribution of the ARID and PHD1 domain region towards efficient demethylation appears to be through interactions of these domains with the nucleosome, as the catalytic efficiency (*k*_max_/*K*_m_^app^) of KDM5C^1-839^ ΔAP relative to wild type is only 3-fold lower on the substrate H3K4me3 peptide (Figure S1A), as opposed to the 9-fold reduction in catalytic efficiency on the substrate nucleosome. As the ARID and PHD1 domains are poorly functionally characterized in KDM5C, we sought to next investigate the features of the nucleosome that they recognize.

### PHD1 domain inhibits KDM5C catalysis

The PHD1 domain of KDM5C has been previously shown to bind to H3K9me3 through peptide pull down [2]. To interrogate the histone binding and specificity of PHD1, we purified the PHD1 domain and quantified binding to histone peptides by nuclear magnetic resonance (NMR) spectroscopy and bio-layer interferometry (BLI). We observe near identical binding between H3 and H3K9me3 tail peptides, indicating no specific binding of PHD1 towards the H3K9me3 modification (Figure S2A). Furthermore, we observe biphasic binding kinetics of PHD1 binding the H3 tail peptide, indicative of a two step binding mechanism (Figure S2B). Upon titration of the H3 tail, large chemical shift changes occur in the two-dimensional heteronuclear single quantum coherence (HSQC) NMR spectrum of a majority of assigned residues in PHD1 (Figure 2A, Figure S2C). The observed affinity of PHD1 towards the H3 tail is surprisingly weak with a dissociation constant of 130 μM, about 100 fold weaker than the affinity of the homologous PHD1 of KDM5A towards the H3 tail (Figure 2B) [25,28]. Despite this difference in affinity, PHD1 of KDM5C retains similar specificity towards the unmodified H3 tail over H3K4 methylated tail peptides as observed in the PHD1 domains of KDM5A/B (Figure 2B).

**Figure 2.**
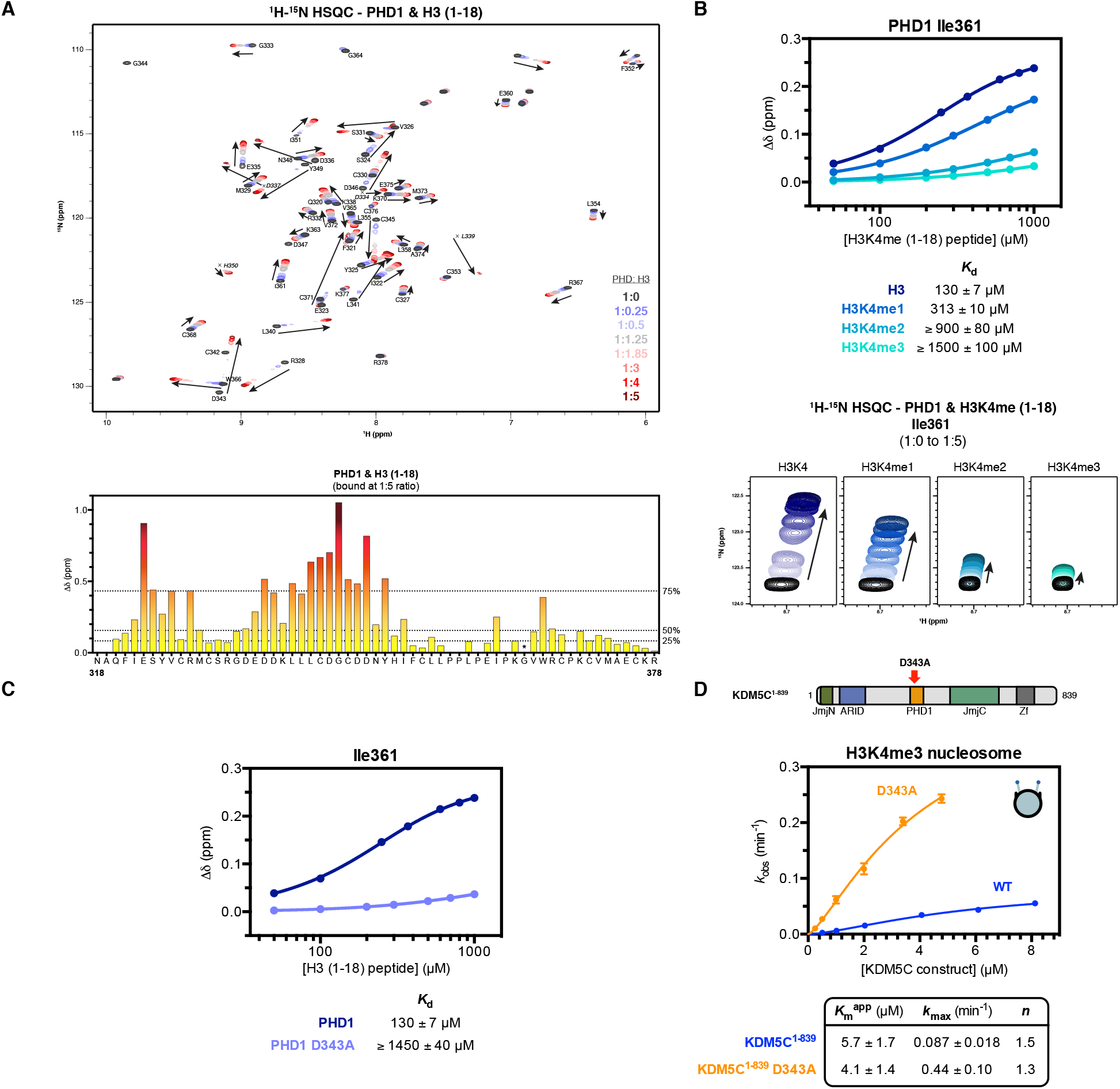
The PHD1 domain of KDM5C preferentially binds the unmodified H3 tail and has an inhibitory role towards nucleosome demethylation. **(A)** 2D ^1^H-^15^N HSQC spectra of PHD1 titrated with increasing amounts of H3 (1-18) peptide with indicated molar ratios (*top*). Backbone assignments of residues in PHD1 are labeled. Corresponding chemical shift change (Δδ) of PHD1 residues upon binding of the H3 (1-18) tail peptide at 1:5 molar ratio (PHD:peptide) (*bottom*). The chemical shift change of G364 (* denoted by asterisk) could not be determined due to broadened chemical shift when bound. Dashed lines indicate 25th, 50th, and 75th percentile rankings, and residues are colored by a gradient from unperturbed (yellow) to significantly perturbed (maroon). Perturbations colored by the gradient and mapped to homologous residues in the structure of KDM5D PHD1 are in Figure S2C. **(B)** 2D ^1^H-^15^N HSQC of I361 in PHD1 upon titration of H3K4me0/1/2/3 (1-18) peptides (*bottom*) with dissociation constants determined from the chemical shift change (Δδ) of I361 with standard error (*top*). Due to incomplete saturation of binding, a lower limit for the dissociation constant is presented for the H3K4me2/3 peptides. Dissociation constants determined from chemical shift changes of several PHD1 residues are in Figure S2H. **(C)** Binding of the H3 (1-18) tail peptide by PHD1 and PHD1 D343A mutant measured by NMR titration HSQC experiments. The chemical shift change (Δδ) of I361 in PHD1 was fit to obtain dissociation constants with standard error. Due to incomplete saturation of binding by the D343A mutant, a lower limit for the dissociation constant is presented. **(D)** Demethylation kinetics of the H3K4me3 substrate nucleosome by wild type and PHD1 mutant KDM5C^1-839^ under single turnover conditions. Observed rates are fit to a cooperative kinetic model, with *n* denoting the Hill coefficient. Wild type kinetic curve replotted from Figure 1C for comparison. Error bars represent SEM of at least three independent experiments (*n* ≥ 3).

In order to investigate the function of PHD1 binding to the H3 tail in KDM5C catalysis, we sought to disrupt the PHD1-H3 interaction through mutagenesis. One of the largest chemical shift perturbations that occurs in PHD1 upon H3 tail binding is at the D343 residue, a residue homologous to D312 in PHD1 of KDM5A where this residue is involved in H3R2 recognition (Figure S2D) [44]. Similarly to PHD1 of KDM5A, we observe a dependence of histone tail binding on recognition of the H3R2 residue by PHD1 of KDM5C (Figure S2E). Like the mutation of D312 in KDM5A, the D343A mutation decreases the affinity of KDM5C PHD1 to the H3 tail at least 10 fold (Figure 2C) [25]. When introduced into the KDM5C^1-839^ enzyme, the D343A mutation does not affect the catalytic rate of H3K4me3 peptide demethylation (Figure S2F). Surprisingly, the D343A PHD1 mutant enzyme demethylates the H3K4me3 nucleosome more rapidly than wild type KDM5C^1-839^, with a ∼5 fold increase of *k*_max_ (Figure 2D). No significant change in nucleosome binding due to the D343A mutation in KDM5C^1-839^ was observed (Figure S2G). This data supports an inhibitory role of the PHD1 domain in nucleosome demethylation by KDM5C. This inhibitory role is in stark contrast to that observed for the PHD1 domain in KDM5A, where the PHD1 domain has a stimulatory role in catalysis [25,28].

### ARID domain contributes to nucleosome demethylation by KDM5C

In contrast to the inhibition of KDM5C demethylation by the PHD1 domain alone, together the ARID and PHD1 domains provide catalytic enhancement on nucleosomes (Figure 1C). We hypothesize that this effect may be due to the ability of the ARID domain to interact with DNA, similarly to the previously demonstrated DNA recognition by the ARID domains of KDM5A/B [13–15]. To test this hypothesis, we interrogated binding of KDM5C^1-839^ towards nucleosomes containing 20 bp flanking DNA on both ends (187 bp nucleosome). Strikingly, we observe a 3-fold gain in affinity towards the 187 bp nucleosome compared to the core (147 bp) nucleosome (Figure 3A), demonstrating that KDM5C is capable of recognizing flanking DNA. KDM5C^1-839^ ΔAP has similar affinity towards both the flanking DNA-containing and core nucleosome (Figure 3B), indicating that the ARID and PHD1 region is responsible for the recognition of flanking DNA.

**Figure 3.**
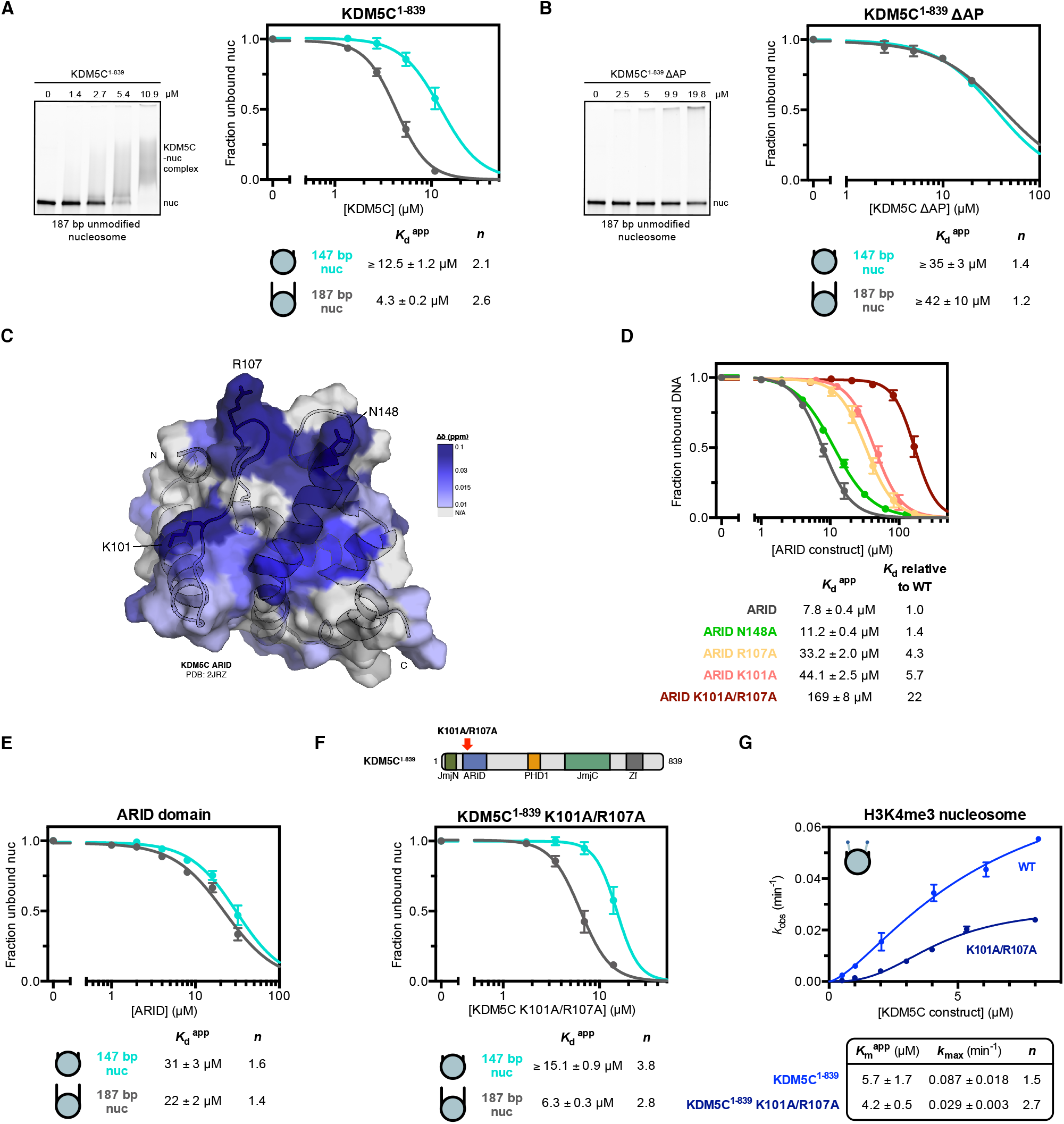
DNA recognition by the ARID domain is needed for nucleosome demethylation but not nucleosome binding by KDM5C. **(A)** Binding of KDM5C^1-839^ to unmodified nucleosomes with and without 20 bp flanking DNA. Representative gel shift of KDM5C binding to the 187 bp nucleosome (*left*). Nucleosome binding curves measured by EMSA and fit to a cooperative binding model to determine apparent dissociation constants (*K*_d_^app^), with *n* denoting the Hill coefficient (*right*). Due to unattainable saturation of binding, a lower limit for the dissociation constant is presented for the unmodified core nucleosome. **(B)** Binding of KDM5C^1-839^ ΔAP to unmodified nucleosomes with and without 20 bp flanking DNA. Representative gel shift of KDM5C ΔAP binding to the 187 bp nucleosome (*left*) and nucleosome binding curves of KDM5C^1-839^ ΔAP (*right*). Due to unattainable saturation of binding, a lower limit for the dissociation constant is presented. **(C)** Chemical shift changes of ARID binding to 20 bp 5’ flanking DNA colored by the gradient and mapped to the KDM5C ARID structure (PDB: 2JRZ) of residues with backbone assignments in the ^1^H-^15^N HSQC spectrum. Significantly perturbed residues are labeled. **(D)** DNA (147 bp 601 core nucleosome positioning sequence) binding by ARID and ARID mutants. Binding curves were measured by EMSA and fit to a cooperative binding model to determine apparent dissociation constants (*K*_d_^app^). **(E)** Nucleosome binding curves of the ARID domain binding to unmodified nucleosomes with and without 20 bp flanking DNA. **(F)** Nucleosome binding curves of ARID mutant KDM5C^1-839^ K101A/R107A binding to unmodified nucleosomes with and without 20 bp flanking DNA. **(G)** Demethylation kinetics of the H3K4me3 core substrate nucleosome by wild type and ARID mutant KDM5C^1-839^ under single turnover conditions. Observed rates are fit to a cooperative kinetic model, with *n* denoting the Hill coefficient. Wild type kinetic curve replotted from Figure 1C for comparison. All error bars represent SEM of at least three independent experiments (*n* ≥ 3).

To further analyze DNA recognition, we purified the KDM5C ARID domain and interrogated its ability to bind the flanking DNA present in the 187 bp nucleosome used in this study. We find that the ARID domain binds the 5’ flanking DNA fragment, with a dissociation constant of 10 μM (Figure S3A). Minimal binding was observed for the 3’ flanking DNA fragment (Figure S3A), suggesting sequence specificity in DNA binding by ARID. We utilized NMR spectroscopy to identify the residues of the ARID domain involved in DNA binding. Previously determined assignments for the ARID domain were reliably transferred to a majority of resonances observed in the ^1^H-^15^N HSQC of ARID, and modest chemical shift changes of select ARID residues were observed upon titration of the 5’ flanking DNA fragment (Figure S3B, Figure S3C) [45]. The perturbed residues localize to a surface on the structure of KDM5C ARID (Figure 3C), with the most notable chemical shift changes at the K101, V105, E106, R107, and N148 residues [45].

We interrogated the contributions of several identified residues, K101, R107, and N148, towards DNA binding through mutagenesis, where we tested binding to the 147 bp 601 core nucleosome positioning sequence (Figure 3D). We find the N148A mutation does not significantly affect DNA binding by ARID, while the K101A and R107A mutations reduce DNA binding by 4-6 fold (Figure 3D). A further 22-fold reduction in DNA binding was observed upon the K101A/R107A double mutation in ARID (Figure 3D), indicating that the K101 and R107 residues are significantly involved in DNA recognition. These residues parallel those identified in the ARID domains of KDM5A/B where the homologous residues, R112 of KDM5A and K119 & R125 of KDM5B, contribute to DNA binding, suggesting conservation of DNA binding residues in the KDM5 family [13,15].

We next interrogated DNA binding by ARID in the context of the 147 bp and 187 bp nucleosomes. We find the ARID domain does not display a strong binding preference for the flanking DNA-containing nucleosome and instead binds both nucleosomes with a similar weak affinity (Figure 3E). The observed binding corresponds to a 3-4 fold reduction in affinity relative to 147 bp non-nucleosomal DNA (Figure 3D, Figure 3E).

We then investigated the function of ARID in the context of the KDM5C enzyme towards nucleosome binding and demethylation by introducing the K101A/R107A double mutation into KDM5C^1-839^. We find that ARID mutant KDM5C^1-839^ retains a similar binding affinity as wild type KDM5C^1-839^ towards both the flanking DNA-containing and core nucleosome (Figure 3F, Figure 3A). This indicates that the ARID domain does not contribute to nucleosome binding or to recognition of flanking DNA by KDM5C, in contrast to our original hypothesis. However, ARID mutant KDM5C^1-839^ has a reduced ability to demethylate the H3K4me3 nucleosome, with a 3-fold reduction in *k*_max_ relative to wild type KDM5C^1-839^ (Figure 3G). Reduced catalysis by the ARID mutant enzyme is only observed on the nucleosome, as the K101A/R107A double mutation does not reduce the catalytic rate of H3K4me3 peptide demethylation (Figure S3D). The similarity of catalytic rates of nucleosome demethylation between ARID mutant KDM5C^1-839^ and KDM5C^1-839^ ΔAP (0.029 min^−1^ and 0.032 min^−1^, respectively) implicates the ARID-DNA interaction as the significant contributor in the ARID and PHD1 region towards catalysis rather than nucleosome recognition (Figure 3G, Figure 1C).

### PHD1 regulates the ability of KDM5C to recognize flanking DNA on the nucleosome

Unlike wild type (Figure 3A) and ARID mutant KDM5C (Figure 3F), KDM5C^1-839^ ΔAP has reduced nucleosome binding and a loss in the ability to discriminate between the 147 bp and 187 bp nucleosome (Figure 3B). To better understand elements of KDM5C that contribute to its ability to bind DNA in the context of the nucleosome, we focused on the linker region between ARID and PHD1. The ARID-PHD1 linker region of KDM5C is the longest among KDM5 family members and contains many basic residues (Figure S4A). This linker region also has low conservation in the KDM5 family and is predicted to be disordered in KDM5C (Figure S4A, Figure S4B). We generated a construct where the linker region (residues 176 to 317) is replaced by a short (GGS)_5_ linker sequence (KDM5C^1-839^ Δlinker) (Figure 4A). KDM5C^1-839^ Δlinker possesses similar catalytic efficiencies as wild type KDM5C^1-839^ on both the H3K4me3 nucleosome and H3K4me3 peptide substrate (Figure S4C, Figure S4D), indicating that the enzyme without the ARID-PHD1 linker is functionally active. We then assessed binding of KDM5C^1-839^ Δlinker to the 147 bp and 187 bp nucleosome and surprisingly did not detect any nucleosome binding (Figure 4A), suggesting that the linker region affects nucleosome and flanking DNA recognition by KDM5C.

**Figure 4.**
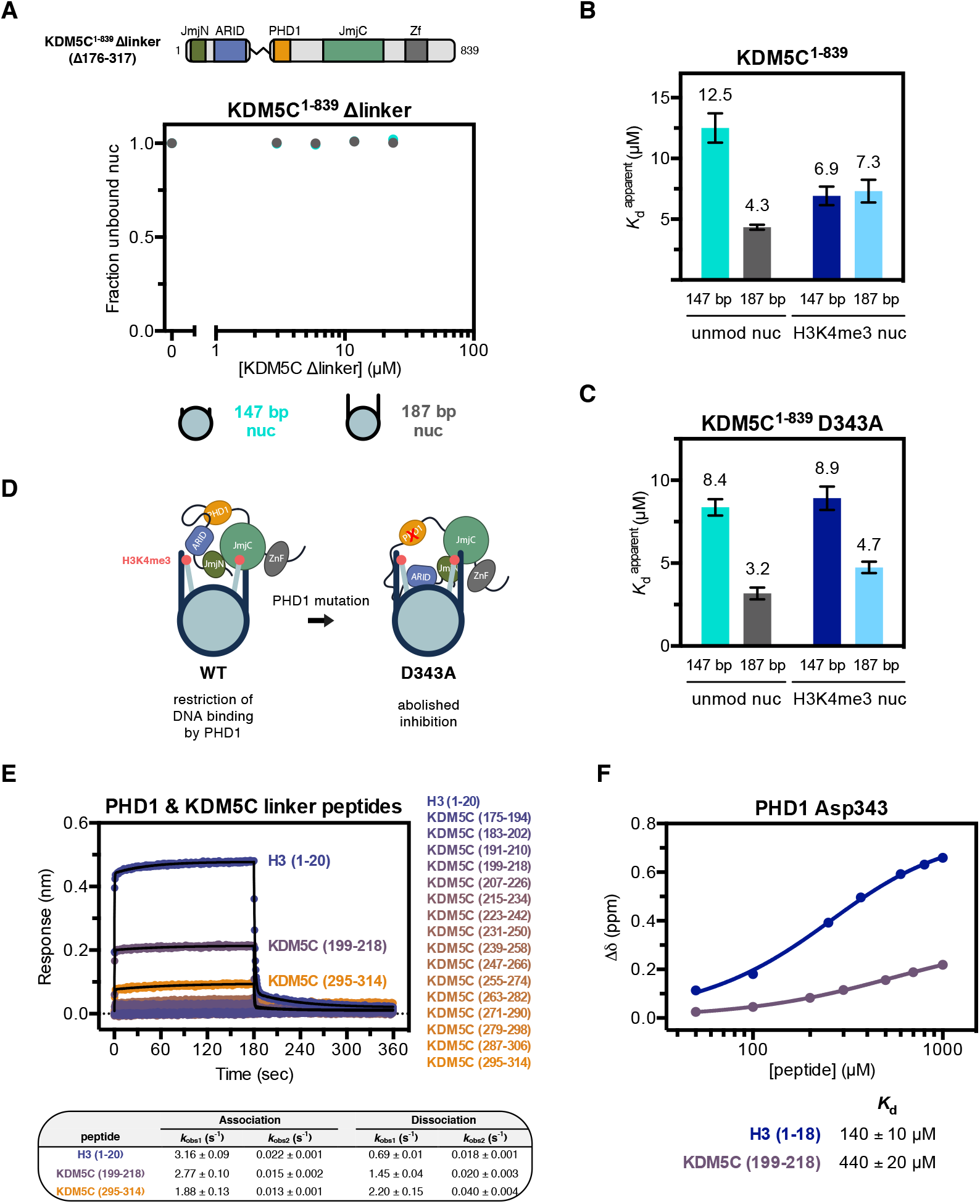
KDM5C recognizes flanking DNA in the absence of H3K4me3 due to regulation by PHD1. **(A)** Binding of KDM5C^1-839^ Δlinker to unmodified nucleosomes with and without 20 bp flanking DNA. Nucleosome binding curves were measured by EMSA. **(B)** Nucleosome binding by KDM5C^1-839^ with apparent dissociation constants (*K*_d_^app^) measured by EMSA and fit to a cooperative binding model (substrate nucleosome binding curves in Figure S4H). Select dissociation constants replotted from Figure 1B and Figure 3A for comparison. Due to unattainable saturation of binding, a lower limit for the dissociation constant is presented for the unmodified core nucleosome. **(C)** Nucleosome binding by PHD1 mutant KDM5C^1-839^ D343A with apparent dissociation constants (*K*_d_^app^) measured by EMSA (binding curves in Figure S4I). **(D)** Model for KDM5C inhibition, where PHD1 prevents flanking DNA recognition in the presence of H3K4me3, and its relief by the PHD1 mutation that disrupts the inhibition. **(E)** Binding kinetic trace of immobilized Avitag-PHD1 binding to H3 (1-20) tail peptide and KDM5C ARID-PHD1 linker fragment 20-mer peptides measured by bio-layer interferometry. Observed rates (*k*_obs_) of association and dissociation by peptides with detectable binding were obtained from fitting kinetic traces to a two phase exponential function. KDM5C linker fragment peptides have acetylated N-termini and amidated C-termini. Identified PHD1-binding KDM5C peptide sequences are KDM5C (199-218): QSVQPSKFNSYGRRAKRLQP and KDM5C (295-314): KEELSHSPEPCTKMTMRLRR. **(F)** Binding of the H3 (1-18) tail peptide and KDM5C (199-218) peptide by PHD1 measured by NMR titration HSQC experiments. The chemical shift change (Δδ) of D343 in PHD1 was fit to obtain dissociation constants with standard error. All error bars represent SEM of at least three independent experiments (*n* ≥ 3).

We next interrogated recognition of flanking DNA on the nucleosome in the presence of the H3K4me3 substrate, as recognition of both could facilitate recruitment of KDM5C to its target sites in euchromatin [29]. Intriguingly, KDM5C^1-839^ has similar binding affinity for both the core and flanking DNA-containing H3K4me3 nucleosome, with *K*_d_^app^ of ∼7 μM, indicating no engagement of flanking DNA in the presence of the H3K4me3 substrate (Figure 4B). This is in contrast to unmodified nucleosome binding, where KDM5C has a clear preference for nucleosomes with flanking DNA (Figure 4B).

Since KDM5C recognizes flanking DNA only in the context of the unmodified nucleosome, we considered the possibility that the ability to engage flanking DNA is coupled to binding of the H3 tail product to the PHD1 domain. To test this model, we interrogated the effect of the PHD1 D343A mutation, which abrogates H3 binding, on the recognition of flanking DNA by KDM5C. We find that PHD1 mutant KDM5C^1-839^ D343A still retains the ∼3-fold affinity gain towards the unmodified 187 bp nucleosome (*K*_d_^app^ = 3.2 μM) compared to the unmodified core nucleosome (*K*_d_^app^ = 8.4 μM) (Figure 4C). In addition, PHD1 mutant KDM5C displays a ∼2 fold affinity gain towards the 187 bp H3K4me3 nucleosome (*K*_d_^app^ = 4.7 μM), relative to the H3K4me3 core nucleosome (*K*_d_^app^ = 8.9 μM) (Figure 4C). Although modest, this improved binding demonstrates that, unlike wild type KDM5C, PHD1 mutant KDM5C can recognize flanking DNA in the presence of the H3K4me3 substrate. This observation lead us to hypothesize that, beyond disruption of H3 tail binding, the D343A mutation may also disrupt intramolecular interactions within the demethylase which restrict the ability of ARID and the ARID-PHD1 linker to interact with DNA (Figure 4D). This PHD1-imposed inhibition model is consistent with the strong catalytic enhancement observed with the PHD1 mutant demethylase under single turnover conditions (Figure 2D).

To further test this model, we examined whether PHD1 is capable of engaging in intramolecular interactions within KDM5C, which could impede its ability to interact with the H3 tail. An intramolecular interaction would necessitate that PHD1 is able to interact with ligands that do not have a free N-terminus, in contrast to typical PHD-H3 interactions [22]. We first tested whether a free N-terminus is required for H3 tail recognition by PHD1 and find that N-terminal acetylation of the H3 tail peptide slightly reduces but does not abrogate binding by PHD1 (Figure S4E), indicating permissibility for recognition of an internal protein sequence. No interaction was detected between PHD1 and the ARID domain (Figure S4F). Using tiled peptides, we then tested binding to peptide fragments of the ARID-PHD1 linker region, each consisting of 20 amino acids with an acetylated N-terminus and amidated C-terminus. PHD1 exhibits most notable binding to the fragment of the ARID-PHD1 linker spanning residues 199-218, which contains a polybasic segment reminiscent of H3 (Figure 4E). Using NMR, titration of PHD1 with the KDM5C (199-218) peptide engages a subset of residues that participate in H3 tail binding, including D343 (Figure S4G, Figure 2A). The affinity of PHD1 towards KDM5C (199-218) is ∼3-fold lower than that of the H3 tail (Figure 4F). These findings indicate PHD1 could interact with the ARID-PHD1 linker within KDM5C, an interaction that can be outcompeted by its H3 tail ligand.

### X-linked intellectual disability mutations alter nucleosome recognition and demethylation by KDM5C

Our proposed regulatory model provides a mechanistic framework for querying the effects of mutations in KDM5C that cause XLID (Figure 5A). Specifically, we sought to investigate the D87G and A388P mutations found at the beginning of ARID and immediately downstream of PHD1, respectively. The D87G mutation, associated with mild intellectual disability, has been demonstrated to have no effect on global H3K4me3 levels *in vivo* [42]. The A388P mutation, associated with moderate intellectual disability, has also been shown to have no effect on global H3K4me3 levels *in vivo* but has been reported to reduce demethylase activity *in vitro* [2,46]. We initially interrogated nucleosome binding by KDM5C^1-839^ D87G and A388P. Strikingly, relative to wild type KDM5C^1-839^, we observe 4-7 fold enhanced binding of the XLID mutants to the unmodified core nucleosome (Figure 5B), suggesting that these mutations enable enhanced nucleosome engagement. The ARID and PHD1 region is required for this enhanced nucleosome binding, as there is no gain in nucleosome affinity due to the A388P mutation when the ARID and PHD1 region is removed (Figure S5A). Importantly, the gain in nucleosome affinity of the XLID mutants is more prominent on the unmodified core nucleosome than the substrate H3K4me3 core nucleosome, resulting in loss of binding specificity towards H3K4me3 by KDM5C due to the D87G and A388P mutations (Figure 5C).

**Figure 5.**
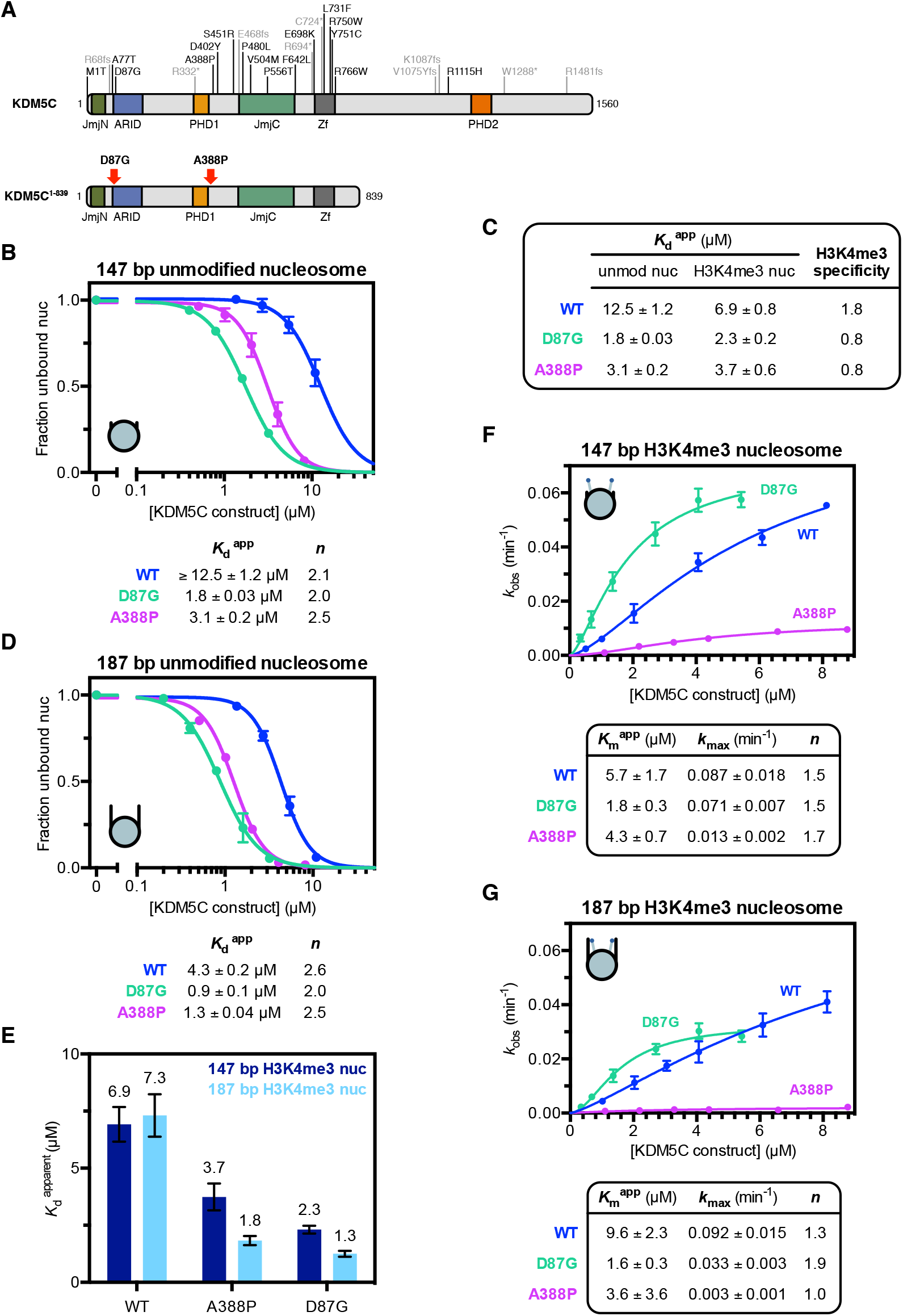
X-linked intellectual disability mutations enhance nucleosome binding by KDM5C and reduce demethylase activity in the presence of flanking DNA. **(A)** XLID mutations found in KDM5C (*top*) and the XLID mutations investigated in this study (*bottom*). **(B)** Unmodified core nucleosome binding by KDM5C^1-839^ wild type (WT), D87G, and A388P. Nucleosome binding was measured by EMSA and curves fit to a cooperative binding model to determine apparent dissociation constants (*K*_d_^app^), with *n* denoting the Hill coefficient. WT binding curve replotted from Figure 3A for comparison. Due to unattainable saturation of binding, a lower limit for the dissociation constant is presented for WT KDM5C binding the unmodified nucleosome. **(C)** Apparent dissociation constants (*K*_d_^app^) of binding by KDM5C^1-839^ WT, D87G, and A388P to unmodified and substrate core nucleosomes and resulting H3K4me3 fold binding specificity. Select dissociation constants are from Figure 1B and Figure 5B for comparison. **(D)** Binding curves of KDM5C^1-839^ WT, D87G, and A388P binding to the unmodified 187 bp nucleosome with 20 bp flanking DNA. WT binding curve replotted from Figure 3A for comparison. **(E)** Binding of KDM5C^1-839^ WT, D87G, and A388P to substrate nucleosomes with and without 20 bp flanking DNA with apparent dissociation constants (*K*_d_^app^) measured by EMSA (binding curves in Figure S5C). Select dissociation constants are replotted from Figure 4B and Figure 5C for comparison. **(F)** Demethylation kinetics of the core substrate nucleosome by KDM5C^1-839^ WT, D87G, and A388P under single turnover conditions. Observed rates are fit to a cooperative kinetic model, with *n* denoting the Hill coefficient. Wild type kinetic curve replotted from Figure 1C for comparison. (G) Demethylation kinetics of the 187 bp substrate nucleosome by KDM5C^1-839^ WT, D87G, and A388P under single turnover conditions. All error bars represent SEM of at least three independent experiments (*n* ≥ 3).

As the XLID mutations cause an overall affinity gain towards both unmodified and substrate nucleosomes, we reasoned that the recognition of the shared common epitope of DNA, rather than the H3 tail, is altered in the mutants. Indeed, relative to wild type KDM5C^1-839^, we observe a similar 3-5 fold gain in affinity by the XLID mutants towards the 187 bp unmodified nucleosome with flanking DNA, with both D87G and A388P mutants converging to a high nucleosome affinity of *K*_d_^app^ ∼1 μM (Figure 5D). As flanking DNA recognition by KDM5C appears to be regulated by PHD1 (Figure 4C), we further interrogated recognition of the 187 bp substrate nucleosome by the D87G and A388P mutants. Both KDM5C^1-839^ D87G and A388P are capable of recognizing flanking DNA in the presence of H3K4me3, with a ∼2 fold gain in affinity towards the 187 bp H3K4me3 nucleosome over the H3K4me3 core nucleosome (Figure 5E). These findings suggest that, similarly to the D343A PHD1 mutation (Figure 4C), the XLID mutations may disrupt the PHD1-mediated inhibition of DNA binding. Our findings are consistent with the model that these XLID mutations are altering the ARID and PHD1 region to relieve the inhibition of DNA binding, enabling unregulated binding to the nucleosome.

We next measured the demethylase activity of KDM5C^1-839^ D87G and A388P towards the H3K4me3 core nucleosome substrate. Despite these XLID mutants sharing similar enhanced nucleosome binding, their effects on nucleosome demethylation differ. The A388P mutation impairs KDM5C catalysis (*k*_max_) by ∼7 fold, while the D87G mutation increases catalytic efficiency (*k*_max_/*K*_m_^app^) ∼3 fold through an enhanced *K*_m_^app^, indicating both nonproductive and productive KDM5C states caused by these mutations (Figure 5F). The reduced demethylase activity caused by the A388P mutation is consistent with previous findings of reduced *in vitro* demethylation, with the 7-fold reduction we observe on nucleosomes exceeding the previously reported 2-fold reduction on substrate peptide [2]. The reduced demethylase activity due to the A388P mutation might be caused by impairment of the composite catalytic domain, as we observe reduced demethylase activity in A388P mutant KDM5C^1-839^ ΔAP (Figure S5B). In contrast, the D87G mutation does not appear to affect the catalytic domain, and instead the improved catalytic efficiency reflects the enhancement in nucleosome binding.

Unlike wild type KDM5C, these XLID mutants recognize flanking DNA in the presence of H3K4me3, prompting us to measure demethylase activity on the 187 bp H3K4me3 nucleosome. Interestingly, while catalysis by the wild type enzyme is only slightly reduced, we find that addition of flanking DNA to the substrate nucleosome results in strong inhibition of catalysis by KDM5C^1-839^ A388P, with a 5-fold reduction in *k*_max_ relative to the core substrate nucleosome (Figure 5G). Addition of flanking DNA also reduces catalysis by KDM5C^1-839^ D87G, although to a lesser degree of ∼2 fold (Figure 5G). Despite lower maximal catalysis (*k*_max_) of the D87G mutant relative to wild type KDM5C^1-839^ in the presence of flanking DNA, the D87G mutant is still ∼2 fold more efficient (*k*_max_/*K*_m_^app^) due to its enhanced nucleosome binding. Regardless, enhanced DNA recognition caused by the XLID mutations results in a reduction in the catalytic rate of H3K4me3 demethylation of nucleosomes with flanking DNA compared to core nucleosomes.

## DISCUSSION

Different reader and regulatory domains within chromatin binding proteins and modifying enzymes influence their activity and substrate specificity by recognizing distinct chromatin states through distinguishing features on the nucleosome and surrounding DNA. Emerging structural studies of chromatin modifying enzymes in complex with nucleosomes have highlighted these multivalent interactions, with increasing observations of interactions with DNA contributing to nucleosome engagement by histone modifying enzymes [47–59]. Despite the unique insertion of the ARID and PHD1 reader domains in the composite catalytic domain, the function of accessory domains within the KDM5 demethylase family has not been explored on nucleosomes. Here, we describe a hierarchy of regulation by these domains by investigating nucleosome recognition and demethylation in KDM5C, a unique member of the KDM5 family involved in regulation of neuronal gene transcription. We find that there are opposing roles of the ARID and PHD1 domains, with DNA recognition by ARID providing a beneficial interaction for nucleosome demethylation and regulation by PHD1 inhibiting nucleosome recognition and demethylation. We further demonstrate that DNA recognition is regulated by the PHD1 domain through its interaction with the ARID-PHD1 linker, allowing for sensing of the H3K4me3 substrate. These regulatory interactions are disrupted by the D87G and A388P XLID mutations adjacent to the ARID and PHD1 domains, resulting in enhanced DNA binding and loss of H3K4me3 specificity. As enhanced flanking DNA recognition by XLID mutants is detrimental to demethylase activity, our findings suggest dysregulation of KDM5C demethylation at euchromatic loci, where this enzyme predominantly functions [29,32].

Our findings of KDM5C nucleosome recognition and demethylation can be best explained by a regulatory model where PHD1 controls DNA recognition (Figure 6A). In the ground state, binding of the ARID-PHD1 linker to PHD1 restricts the ability of the enzyme to interact with DNA, attenuating catalysis (state I). Release of the PHD1-imposed constraint on the ARID-PHD1 linker and ARID domain enables improved interaction with DNA, leading to faster catalysis (state II). In our experiments, the D343A PHD1 mutation was used as a mechanistic probe to release the PHD1-imposed restriction on DNA binding. In the context of chromatin, this release of inhibition could be achieved through binding of the H3 tail to PHD1, allowing for the regulation of demethylation by the surrounding chromatin environment. Formation of the demethylated H3 product, and its binding to PHD1, further reinforces an interaction of KDM5C with chromatin by enabling flanking DNA recognition, possibly through the ARID-PHD1 linker region (state III). Alternatively, the PHD1 domain could act directly on the catalytic domains to impair productive substrate nucleosome engagement.

**Figure 6.**
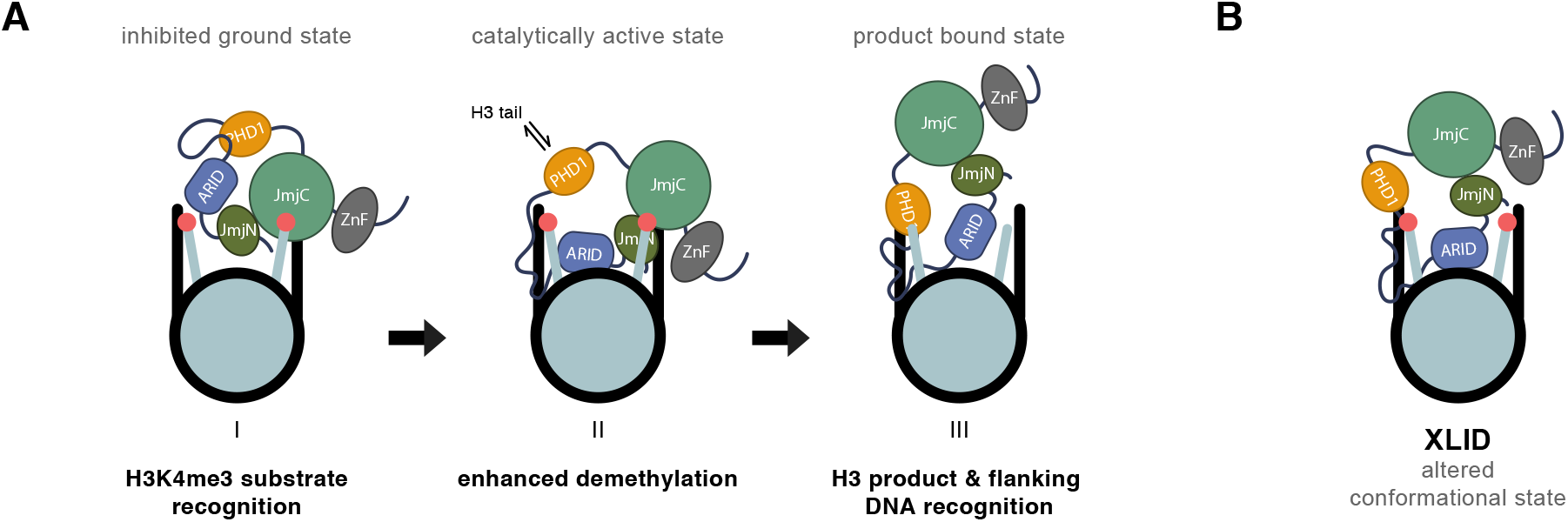
Model of KDM5C regulation by the ARID-linker-PHD1 region and KDM5C dysregulation by XLID mutations. **(A)** KDM5C recognizes H3K4me3 and binds to substrate nucleosomes through the catalytic domain (*pre-catalytic and inhibited ground state I)*. DNA binding in the presence of H3K4me3 is attenuated due to an inhibitory role of PHD1 on DNA recognition. During demethylation, ARID makes transient interactions with nucleosomal DNA to orient the catalytic domain towards the H3K4me3 tail for efficient demethylation. H3 tail binding to PHD1 releases the PHD1 interaction constraining the ARID-PHD1 linker and ARID domain, enabling ARID interactions with DNA to further enhance demethylation (*catalytically active state II)*. After demethylation, binding of the product H3 tail to PHD1 enables flanking DNA binding by the ARID-PHD1 linker region (*post-catalytic and product bound state III)*. **(B)** Proposed altered conformational state of the ARID and PHD1 region in KDM5C due to XLID mutations in this region disrupting hypothesized intramolecular interactions.

Intriguingly, we observe cooperativity (Hill coefficients > 1) in nucleosome binding and demethylation (Figure 1C, Figure S1B). In addition, cooperativity occurs in peptide demethylation by wild type KDM5C^1-839^ but not by KDM5C^1-839^ ΔAP under single turnover conditions (Figure S1A), suggesting that cooperativity might arise both from the state of KDM5C and from the nucleosome possessing two H3 tails.

Our finding of the beneficial role of the ARID domain towards KDM5C catalysis on nucleosomes can be rationalized by favorable transient interactions of the ARID domain with nucleosomal DNA to better orient the catalytic domains for demethylation and could make the substrate H3K4me3 more accessible through disrupting histone tail-DNA interactions [60–63]. This is supported by the previous observation that the ARID domain of KDM5C is required for its demethylase activity *in vivo* but not for its chromatin association [11]. This role of the ARID domain in productive nucleosome demethylation may be conserved within the KDM5 family, as the ARID domain has also been found to be required for *in vivo* demethylation by KDM5A/B and the Drosophila KDM5 homolog Lid [4,10,12,13]. The ARID domain may be required for nucleosome demethylation in order to displace the H3K4me3 tail from interacting with DNA, making it accessible for engagement by the catalytic domain. This histone tail displacement function has been proposed for DNA binding reader domain modules and for the LSD1/CoREST complex, where the SANT2 domain interacts with nucleosomal DNA to displace the H3 tail for engagement by the LSD1 active site [55,63–65].

In contrast to the beneficial role of the ARID domain, we observe an unexpected inhibitory role of PHD1 towards KDM5C demethylation on nucleosomes. This finding suggests differential regulation by PHD1 in the KDM5 family, as PHD1 binding has a stimulatory role towards *in vitro* demethylation in KDM5A/B, and PHD1 has been previously shown to be required for demethylase activity *in vivo* for KDM5B and Lid [4,10,25,26,28]. Our data suggests this inhibitory role is mediated by the ability of PHD1 to inhibit KDM5C’s engagement of DNA on the nucleosome (Figure 6A). With weak affinity and indifference for a free N-terminus (Figure S4E), ligand recognition by PHD1 in KDM5C is strikingly different from that observed for the PHD1 domains in KDM5A and KDM5B. While further work is needed to identify how PHD1 restricts DNA binding, our findings indicate that this could be achieved through an interaction between PHD1 and the unstructured ARID-PHD1 linker region, possibly mediated by the basic residues within the linker. This unique ARID-PHD1 linker (Figure S4A) may contribute to distinct regulation by PHD1 in KDM5C. Although we are unable to directly test the effect of H3 tail binding to PHD1 on DNA recognition due to the low affinity regime, we hypothesize that the resulting binding could release inhibition, allowing for the regulation of KDM5C activity by different chromatin environments. As a consequence, H3 tail binding by PHD1 might stimulate demethylation, as observed upon PHD1 binding in KDM5A/B, through a mechanistically distinct relief of negative regulation in KDM5C (Figure 6A).

Unlike the ARID domain, whose DNA recognition is needed for nucleosome demethylation but not nucleosome binding, the ARID-PHD1 linker region contributes towards nucleosome binding but does not appear to contribute to demethylation by KDM5C. Surprisingly, we observe diminished nucleosome binding upon deletion of the ARID-PHD1 linker as opposed to a ∼3-fold decrease in nucleosome binding upon deletion of the entire ARID and PHD1 region. While the molecular basis for these effects requires further studies, this observed discrepancy could result from the ARID and PHD1 domains affecting nucleosome binding by the catalytic and zinc finger domains of KDM5C. Our findings add to the reports of intrinsically disordered regions as functional elements within chromatin binding proteins [66–69].

Unexpectedly, KDM5C recognizes flanking DNA around the nucleosome in the presence of the unmodified H3 tail but not in the presence of the H3K4me3 substrate. Linker DNA recognition may serve to retain KDM5C at its target promoter and enhancer sites within open chromatin after demethylation. It may also enable processive demethylation of adjacent nucleosomes in euchromatin by KDM5C. Interestingly, the recognition of linker DNA has been observed in the mechanistically unrelated H3K4me1/2 histone demethylase LSD1/KDM1A, where demethylase activity is in contrast stimulated by linker DNA [47,70]. The H3K36me1/2 demethylase KDM2A is also capable of recognizing linker DNA, where it is specifically recruited to unmethylated CpG islands at gene promoters through its ZF-CxxC domain [71,72]. These findings suggest that recognition of the chromatin state with accessible linker DNA may be utilized by histone modifying enzymes that function on euchromatin. While the sequence specificity of linker DNA recognition requires further investigation, it is evident that the sensing of the H3K4me3 substrate tail by KDM5C is preferred over recognition of linker DNA, a feature accessible in open chromatin. This observed hierarchy, coupled with KDM5C’s overall weak affinity towards nucleosomes and dampened demethylase activity due to regulation by PHD1, suggests tunable demethylation by KDM5C. Thus, this multi-domain regulation might serve to establish H3K4me3 surveillance through KDM5C-catalyzed demethylation, which is well suited for the physiological role of this enzyme in fine tuning gene expression through H3K4me3 demethylation at enhancers and promoters of genes, as well as its role in genome surveillance by preventing activation of non-neuronal genes in adult neurons [29,32].

Our findings show that the regulation of DNA recognition by KDM5C is disrupted by the D87G and A388P XLID mutations adjacent to the ARID and PHD1 domains, such that nucleosome binding is significantly enhanced, H3K4me3 specificity is lost, and demethylase activity is sensitized to inhibition by linker DNA. The location of these mutations lends support to our model, where the XLID mutations relieve the inhibition of DNA recognition, enabling enhanced nucleosome binding irrespective of the methylation status of the nucleosome, by altering the conformational state of the ARID and PHD1 region (Figure 6B). Beyond disruption of histone demethylase activity, our findings suggest an additional mechanism of dysregulation of KDM5C in XLID, that of enhanced nonproductive chromatin engagement and differential dysregulation of demethylation at different loci depending on the accessibility of linker DNA. Despite the reduced *in vitro* activity of KDM5C due to the A388P mutation, global H3K4me3 levels are unaffected with human KDM5C A388P *in vivo* [2,46]. In contrast, increased global H3K4me3 levels have been observed in a Drosophila intellectual disability model with A512P mutant Lid, signifying that further work is needed to profile H3K4me3 levels at genomic target regions affected by XLID mutations in human KDM5C [73]. Furthermore, we observe that the demethylase activity of KDM5C D87G varies relative to wild type depending on the presence of flanking DNA, which might account for the unaffected global H3K4me3 levels previously observed with this D87G mutation [42]. Our findings suggest that the chromatin environment, in particular the presence of accessible linker DNA, could govern altered demethylation and nonproductive chromatin recognition by KDM5C in XLID. Euchromatin-specific dysregulation of KDM5C demethylation might account for the hard-to-reconcile discrepancies between reported *in vitro* demethylase activities of KDM5C XLID mutants and their effect on global H3K4me3 levels.

## MATERIALS AND METHODS

### Generation of KDM5C constructs

Human KDM5C gene was obtained from Harvard PlasmID (HsCD00337804) and Q175 was inserted to obtain the canonical isoform (NP_004178.2). KDM5C residues 1 to 839 were cloned into a pET28b His-Smt3 vector to produce 6xHis-SUMO-KDM5C and was mutated by site-directed mutagenesis for point mutants. The KDM5C^1-839^ ΔAP construct was cloned by replacing residues 83-378 with a 4xGly linker. The KDM5C^1-839^ Δlinker construct was cloned by replacing residues 176-317 with a (GGS)_5_ linker.

### Purification of KDM5C constructs

Recombinant His-tagged SUMO-KDM5C constructs were expressed in BL21(DE3) *E. coli* in LB media containing 50 μM ZnCl_2_ and 100 μM FeCl_3_ through induction at OD_600_ ∼0.6 using 100 μM IPTG followed by expression at 18 °C overnight. Collected cells were resuspended in 50 mM HEPES pH 8, 500 mM KCl, 1 mM BME, 5 mM imidazole, and 1 mM PMSF, supplemented with EDTA-free Pierce protease inhibitor tablets (Thermo Fisher Scientific) and benzonase, and lysed by microfluidizer. Lysate was clarified with ultracentrifugation and the recovered supernatant was then purified by TALON metal affinity resin (total contact time under 2 hrs) at 4 °C. The His-SUMO tag was then cleaved by SenP1 during overnight dialysis at 4 °C in 50 mM HEPES pH 7.5, 150 mM KCl, and 5 mM BME. KDM5C constructs were then purified by anion exchange (MonoQ, GE Healthcare) and subsequent size exclusion (Superdex 200, GE Healthcare) chromatography in 50 mM HEPES pH 7.5 and 150 mM KCl. Fractions were concentrated and aliquots snap frozen in liquid nitrogen for storage at −80 °C.

### Nucleosomes and DNA

Recombinant human 5’ biotinylated unmodified 147 bp mononucleosomes (16-0006), unmodified 187 bp mononucleosomes (16-2104), 5’ biotinylated H3K4me3 147 bp mononucleosomes (16-0316), and 5’ biotinylated H3K4me3 187 bp mononucleosomes (16-2316) were purchased from Epicypher, Inc., in addition to biotinylated 147 bp 601 sequence DNA (18-005). 187 bp nucleosomes contain the 20 bp sequences 5’ GGACCCTATACGCGGCCGCC and GCCGGTCGCGAACAGCGACC 3’ flanking the core 601 positioning sequence. 20 bp flanking DNA duplex fragments were synthesized by Integrated DNA Technologies, Inc. For use in binding and kinetic assays, stock nucleosomes were buffer exchanged into corresponding assay buffer using a Zeba micro spin desalting column (Thermo Scientific).

### Nucleosome and DNA binding assays

Nucleosome and DNA binding was assessed by EMSA. 100 nM nucleosomes (0.5 pmol) and various concentrations of KDM5C were incubated in binding buffer (50 mM HEPES pH 7.5, 50 mM KCl, 1mM BME, 0.01% Tween-20, 0.01% BSA, 5% sucrose) for 1 hr on ice prior to analysis by native 7.5% PAGE. For DNA binding, 100 nM 147 bp 601 sequence DNA or 500 nM 20 bp linker DNA fragments were incubated with various concentrations of ARID. Samples were separated using pre-run gels by electrophoresis in 1xTris-Glycine buffer at 100V for 2 hrs at 4 °C, stained using SYBR Gold for DNA visualization, and imaged using the ChemiDoc imaging system (Bio-Rad Laboratories). Bands were quantified using Bio-Rad Image Lab software to determine the fraction of unbound nucleosome to calculate apparent dissociation constants by fitting to the cooperative binding equation Y=(X^n)/(K_d_^n + X^n), where X is the concentration of KDM5C, n is the Hill coefficient, and K_d_ is the concentration of KDM5C at which nucleosomes are half bound.

### Single turnover nucleosome demethylation kinetics

The demethylation of biotinylated H3K4me3 nucleosome was monitored under single turnover conditions (>10 fold excess of KDM5C over substrate) through the detection of H3K4me1/2 product nucleosome formation over time by TR-FRET of an anti-H3K4me1/2 donor with an anti-biotin acceptor reagent. Various concentrations of KDM5C were reacted with 25 nM 5’ biotinylated H3K4me3 nucleosome in 50 mM HEPES pH 7.5, 50 mM KCl, 0.01% Tween-20, 0.01% BSA, 50 μM alpha-ketoglutarate, 50 μM ammonium iron(II) sulfate, and 500 μM ascorbic acid at room temperature. 5 μL time points were taken and quenched with 1.33 mM EDTA then brought to 20 μL final volume for detection using 1 nM LANCE Ultra Europium anti-H3K4me1/2 antibody (TRF0402, PerkinElmer) and 50 nM LANCE Ultra Ulight-Streptavidin (TRF0102, PerkinElmer) in 0.5X LANCE detection buffer. Detection reagents were incubated with reaction time points for 2 hours at room temperature in 384 well white microplates (PerkinElmer OptiPlate-384) then TR-FRET emission at 665 nm and 615 nm by 320 nm excitation with 50 μs delay and 100 μs integration time was measured using a Molecular Devices SpectraMax M5e plate reader. TR-FRET was calculated as the 665/615 nm emission ratio and kinetic curves were fit to a single exponential function to determine *k*_obs_ of demethylation. *k*_obs_ parameters were then plotted as a function of KDM5C concentration and fit to the sigmoidal kinetic equation Y=k_max_*X^n/(K_half_^n + X^n) using GraphPad Prism to determine *k*_max_ and *K*_m_^app^ parameters of demethylation.

### Purification of PHD1 for NMR

PHD1 (KDM5C residues 318-378) was cloned into a pET28b His-Smt3 vector to express recombinant 6xHis-SUMO-PHD1 in BL21(DE3) *E. coli* in metal supplemented M9 minimal medium containing ^15^NH_4_Cl (Cambridge Isotope Laboratories). ^13^C-glucose (Cambridge Isotope Laboratories) was used in medium for expression of ^15^N, ^13^C-labeled PHD1. Expression was induced at OD_600_ ∼0.6 using 1 mM IPTG for expression at 18 °C overnight. Collected cells were resuspended in 50 mM HEPES pH 8, 500 mM KCl, 5 mM BME, 10 mM imidazole, and 1 mM PMSF, supplemented with benzonase, and lysed by sonication. Lysate was clarified with ultracentrifugation and the recovered supernatant was then purified by Ni-NTA affinity resin. The His-SUMO tag was then cleaved by SenP1 during overnight dialysis at 4 °C in 50 mM HEPES pH 7.5, 150 mM KCl, 50 μM ZnCl_2_ and 10 mM BME. Cleaved His-SUMO tag and SenP1 was captured by passing through Ni-NTA affinity resin and flow-through was then purified by anion exchange (MonoQ) chromatography in starting buffer of 50 mM HEPES pH 7.5, 150 mM KCl, 50 μM ZnCl2 and 10 mM BME. Flow-through MonoQ fractions containing PHD1 were concentrated and aliquots snap frozen in liquid nitrogen for storage at −80 °C.

### PHD1 NMR and histone peptide NMR titrations

For backbone assignment of KDM5C PHD1, 400 μM ^15^N, ^13^C-labeled PHD1 in 50 mM HEPES pH 7.5, 50 mM KCl, 5 mM BME, 50 μM ZnCl_2_, and 5% D_2_O was used to perform 3D triple-resonance CBCA(CO)NH and CBCANH experiments at 298K using a 500 MHz Bruker spectrometer equipped with a cryoprobe. Triple-resonance experiments were also performed using 400 μM ^15^N, ^13^C-labeled PHD1 bound to 2 mM H3 (1-18) peptide (1:5 ratio) to assign broadened backbone residues in apo spectra. 3D spectra were processed using NMRPipe then analyzed and assigned using CcpNMR Analysis. Out of 56 assignable residues, 54 in apo PHD1 and 53 residues in H3 bound PHD1 were assigned.

For 2D ^1^H-^15^N HSQC spectra of KDM5C PHD1, 200 μM ^15^N-labeled PHD1 in 50 mM HEPES pH 7.5, 50 mM KCl, 5 mM BME, 50 μM ZnCl_2_, and 5% D_2_O was used to obtain 2D spectra at 298K using a 800 MHz Bruker spectrometer equipped with a cryoprobe. Chemical shift perturbation experiments were performed by obtaining HSQC spectra with increasing concentrations of histone tail peptides (GenScript) up to 1:5 molar ratio of PHD1:peptide. Data were processed using Bruker TopSpin and analyzed using CcpNMR Analysis. Chemical shifts were scaled and calculated as Δδ = sqrt(((ΔδH)^2+(ΔδN/5)^2) / 2). Chemical shift values were then plotted as a function of histone peptide concentration and fit to the quadratic binding equation Y=((X+P_T_+K_d_)-sqrt((X+P_T_+K_d_)^2-4*P_T_*X))*(Y_max_-Y_min_)/(2*P_T_), where X is the concentration of peptide and P_T_ is the concentration of PHD1, using GraphPad Prism to determine K_d_ values.

### Purification of ARID for NMR

ARID (KDM5C residues 73-188) was cloned into a pET28b His-Smt3 vector to express recombinant 6xHis-SUMO-ARID in BL21(DE3) *E. coli* in metal supplemented M9 minimal medium containing ^15^NH_4_Cl. Expression was induced at OD_600_ ∼0.6 using 1 mM IPTG for expression at 18 °C overnight. Collected cells were resuspended in 50 mM HEPES pH 8, 500 mM KCl, 1 mM BME, 10 mM imidazole, and 1 mM PMSF, supplemented with EDTA-free Pierce protease inhibitor tablets and benzonase, and lysed by microfluidizer. Lysate was clarified with ultracentrifugation and the recovered supernatant was then purified by Ni-NTA affinity resin. The His-SUMO tag was then cleaved by SenP1 during overnight dialysis at 4 °C in 50 mM HEPES pH 7.5, 500 mM KCl, and 5 mM BME. Cleaved His-SUMO tag and SenP1 was captured by passing through Ni-NTA affinity resin and flow-through was then purified by size exclusion (Superdex 75, GE Healthcare) chromatography in 50 mM HEPES pH 7, 150 mM KCl, and 5 mM BME. Fractions were buffer exchanged into 50 mM HEPES pH 7, 50 mM KCl, and 5 mM BME then concentrated and aliquots snap frozen in liquid nitrogen for storage at −80 °C.

### ARID and DNA NMR titration

For 2D ^1^H-^15^N HSQC spectra of KDM5C ARID, 100 μM ^15^N-labeled ARID in 50 mM HEPES pH 7, 50 mM KCl, 5 mM BME, and 5% D_2_O was used to obtain 2D spectra at 298K using a 800 MHz Bruker spectrometer equipped with a cryoprobe. Chemical-shift perturbation experiments were performed by obtaining HSQC spectra with increasing concentrations of the 5’ linker DNA 20 bp fragment up to 1:1 molar ratio of ARID:DNA. For the PHD1 titration experiment, 50 μM ^15^N-labeled ARID in 50 mM HEPES pH 7, 50 mM KCl, 5 mM BME, 50 μM ZnCl_2_, and 5% D_2_O was used with increasing concentrations of PHD1 up to 1:3 molar ratio of ARID:PHD1. Data were processed using Bruker TopSpin and analyzed using CcpNMR Analysis. Chemical shifts were scaled and calculated as Δδ = sqrt(((ΔδH)^2+(ΔδN/5)^2) / 2). Previously determined assignments (BMRB: 15348) were transferred to a majority of resonances observed in the HSQC spectra of ARID [45].

### Purification of ARID mutants

Recombinant His-tagged SUMO-ARID mutants were expressed in BL21(DE3) *E. coli* in 2xTY media through induction at OD_600_ ∼0.6 using 1 mM IPTG followed by expression at 18 °C overnight. Collected cells were resuspended in 50 mM HEPES pH 8, 500 mM KCl, 1 mM BME, 10 mM imidazole, and 1 mM PMSF, supplemented with benzonase, and lysed by sonication. Lysate was clarified with centrifugation and the recovered supernatant was then purified by Ni-NTA affinity resin. The His-SUMO tag was then cleaved by SenP1 for 2 hours at 4 °C in 50 mM HEPES pH 7, 500 mM KCl, and 5 mM BME. Cleaved His-SUMO tag and SenP1 was captured by passing through Ni-NTA affinity resin. The flow-through was buffer exchanged into 50 mM HEPES pH 7, 50 mM KCl, and 5 mM BME then concentrated and aliquots snap frozen in liquid nitrogen for storage at −80 °C.

### Single turnover peptide demethylation kinetics

The demethylation of biotinylated H3K4me3 peptide was monitored under single turnover conditions (>10 fold excess of KDM5C over substrate) through the detection of H3K4me3 substrate loss over time by TR-FRET of an anti-rabbit IgG donor, recognizing an anti-H3K4me3 rabbit antibody, with an anti-biotin acceptor reagent. Various concentrations of KDM5C were reacted with 25 nM H3K4me3 (1-21)-biotin peptide (AS-64357, AnaSpec) in 50 mM HEPES pH 7.5, 50 mM KCl, 0.01% Tween-20, 0.01% BSA, 50 μM alpha-ketoglutarate, 50 μM ammonium iron(II) sulfate, and 500 μM ascorbic acid at room temperature. 2.5 μL time points were taken and quenched with 2 mM EDTA then brought to 20 μL final volume for detection using 1:500 dilution anti-H3K4me3 antibody (05-745R, EMD Millipore), 1 nM LANCE Ultra Europium anti-rabbit IgG antibody (PerkinElmer AD0082), and 50 nM LANCE Ultra Ulight-Streptavidin (PerkinElmer TRF0102) in 0.5X LANCE detection buffer. Detection reagents were added stepwise with 30 min incubation of anti-H3K4me3 antibody and Ulight-Streptavidin with reaction time points followed by 1 hr incubation with Europium anti-rabbit antibody in 384 well white microplates (PerkinElmer OptiPlate-384). TR-FRET emission at 665 nm and at 615 nm by 320 nm excitation with 50 μs delay and 100 μs integration time was measured using a Molecular Devices SpectraMax M5e plate reader. TR-FRET was calculated as the 665/615 nm emission ratio then subject to normalization to H3K4me3 substrate signal before demethylation. Kinetic curves were fit to a single exponential function, with the plateau set to nonspecific background of H3K4me2 product detection, to determine *k*_obs_ of the H3K4me3 demethylation step. *k*_obs_ parameters were then plotted as a function of KDM5C concentration and fit to the sigmoidal kinetic equation Y=k_max_*X^n/(K_half_^n + X^n) using GraphPad Prism to determine *k*_max_ and *K*_m_’ parameters of demethylation.

### Multiple turnover peptide demethylation kinetics

A fluorescence-based enzyme coupled assay was used to detect the formaldehyde product of demethylation of H3K4me3 peptide under multiple turnover conditions (excess of substrate peptide over KDM5C). Various concentrations of H3K4me3 (1-21) substrate peptide (GenScript) were added with 1mM alpha-ketoglutarate to initiate demethylation by ∼1 μM KDM5C in 50 mM HEPES pH 7.5, 50 mM KCl, 50 μM ammonium iron(II) sulfate, 2 mM ascorbic acid, 2 mM NAD+, and 0.05 U formaldehyde dehydrogenase (Sigma-Aldrich) at room temperature. Upon initiation, fluorescence (350 nm excitation, 460 nm emission) was measured in 20 sec intervals over 30 min using a Molecular Devices SpectraMax M5e plate reader. NADH standards were used to convert fluorescence to the rate of product concentration formed. Initial rates of the first 3 min of demethylation were plotted as a function of substrate concentration and fit to the tight-binding quadratic velocity equation Y=V_max_*((X+E_T_+K_m_)-sqrt((X+E_T_+K_m_)^2-4*E_T_*X))/(2*E_T_) using GraphPad Prism to determine Michaelis-Menten kinetic parameters of demethylation.

### Histone peptide binding kinetics

Bio-layer interferometry was used to measure binding kinetics of histone peptides to biotinylated Avitag-PHD1. Avitag followed by a linker was inserted into pET28b His-Smt3-PHD1^318-378^ to generate recombinant endogenously biotinylated 6xHis-SUMO-Avitag-(GS)_2_-PHD1 through coexpression with BirA in BL21(DE3) *E. coli* in 2xTY media containing 50 μM ZnCl_2_ and 50 μM biotin. Expression was induced at OD_600_ ∼0.7 using 0.4 mM IPTG for expression at 18 °C overnight. Collected cells were resuspended in 50 mM HEPES pH 8, 500 mM KCl, 5 mM BME, 10 mM imidazole, 50 μM biotin, and 1 mM PMSF, supplemented with benzonase, and lysed by sonication. Lysate was clarified with ultracentrifugation and the recovered supernatant was then purified by Ni-NTA affinity resin. The His-SUMO tag was then cleaved by SenP1 during overnight dialysis at 4 °C in 50 mM HEPES pH 8, 150 mM KCl, 50 μM ZnCl_2_ and 10 mM BME. Cleaved His-SUMO tag and SenP1 was captured by passing through Ni-NTA affinity resin and flow-through was then purified by anion exchange (MonoQ) chromatography in starting buffer of 50 mM HEPES pH 8, 150 mM KCl, 50 μM ZnCl_2_ and 10 mM BME. Flow-through MonoQ fractions containing Avitag-PHD1 were analyzed by western blotting to identify biotinylated fractions, which were then concentrated and aliquots snap frozen in liquid nitrogen for storage at −80 °C. Using the Octet Red384 system (ForteBio) at 1000 rpm and 25 °C, 100 nM Avitag-PHD1 was loaded onto streptavidin biosensors (ForteBio) for 10 min in assay buffer (50 mM HEPES pH 8, 50 mM KCl, 50 μM ZnCl_2_, 5 mM BME, and 0.05% Tween-20) followed by 120 sec baseline then association and dissociation of 100 μM peptide (GenScript) in assay buffer. Data were processed by subtracting a single reference experiment of loaded Avitag-PHD1 without peptide. A two phase exponential function was used to fit the biphasic kinetic data using Origin software. For the ARID-PHD1 linker peptides tested to bind PHD1, the KDM5C (263-282) and KDM5C (271-290) peptides were found to nonspecifically associate with biosensors, and loaded biosensors were associated with these peptides prior to obtaining their binding traces.

## ACKNOWLEDGMENTS

We thank Barbara Panning, Geeta Narlikar, John Gross, Daniele Canzio, Ryan Tibble, Cynthia Chio, and members of the Fujimori laboratory for helpful discussions and guidance.

## FUNDING AND ADDITIONAL INFORMATION

This work was supported by the UCSF Discovery Fellows program and National Science Foundation Graduate Research Fellowship to F. S. U. and by the National Institutes of Health (R01 GM114044, R01 GM114044-03S1, and R01 CA250459) to D. G. F. The content is solely the responsibility of the authors and does not necessarily represent the official views of the National Institutes of Health.

## SUPPLEMENTAL INFORMATION

This article contains supporting information.

## ABBREVIATIONS

XLID: (X-linked intellectual disability)
KDM: (lysine demethylase)
H3: (histone 3)
H3K4: (histone 3 lysine 4)
H3K4me1: (monomethylated lysine 4 of histone 3)
H3K4me2: (dimethylated lysine 4 of histone 3)
H3K4me3: (trimethylated lysine 4 of histone 3)
PHD: (plant homeodomain)
ARID: (AT-rich interaction domain)
JmjN: (Jumonji N domain)
JmjC: (Jumonji C domain)
ZnF: (zinc finger domain)
KDM5C^1-839^ ΔAP: (KDM5C protein containing residues 1 to 839 with truncation of the ARID and PHD1 region)
KDM5C^1-839^ Δlinker: (KDM5C protein containing residues 1 to 839 with truncation of the linker region between ARID and PHD1)
HSQC: (heteronuclear single quantum coherence)
EMSA: (electrophoretic mobility shift assay)
TR-FRET: (time-resolved fluorescence resonance energy transfer)
nuc: (nucleosome)

## SUPPLEMENTAL INFORMATION

**Figure S1.**
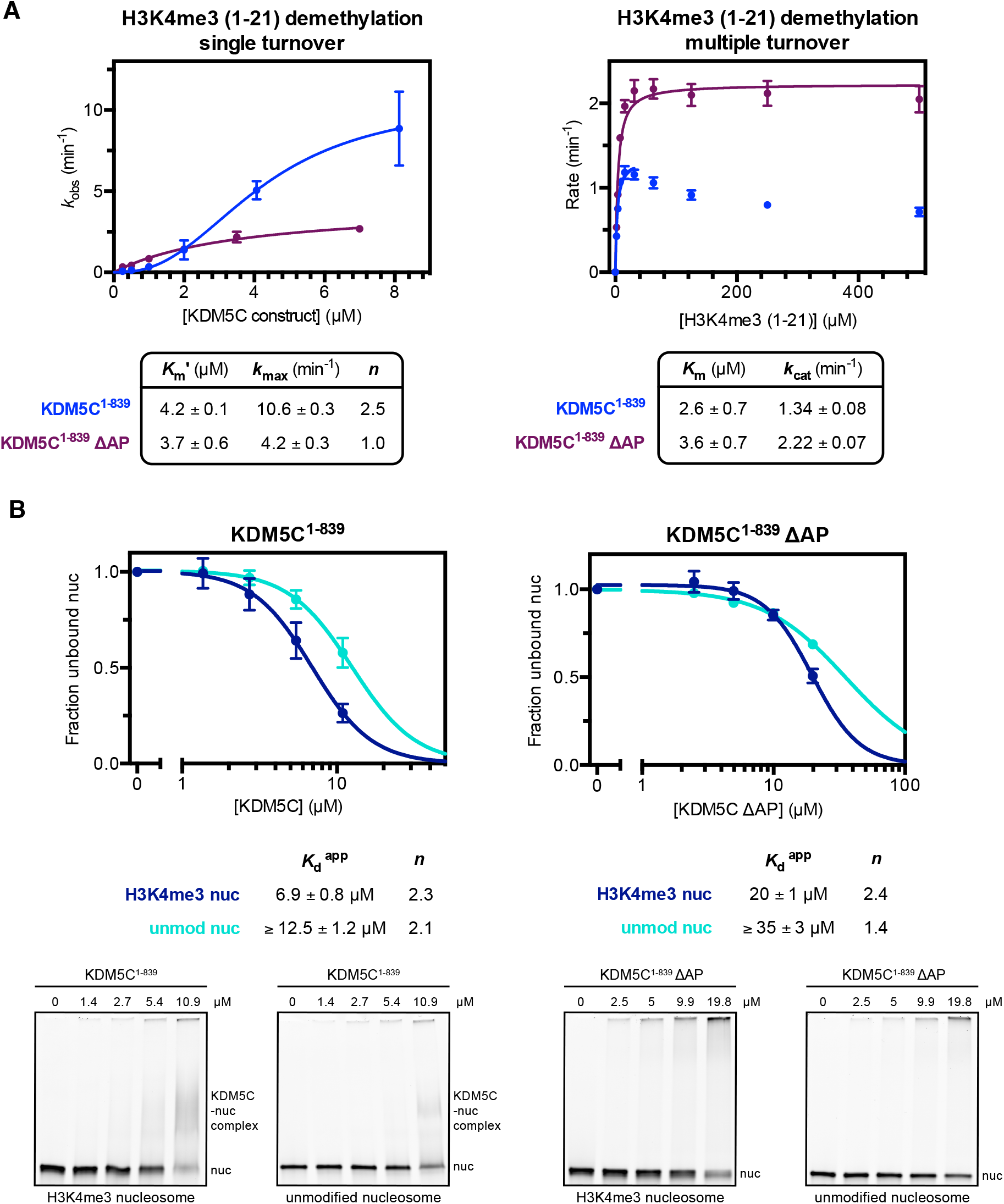
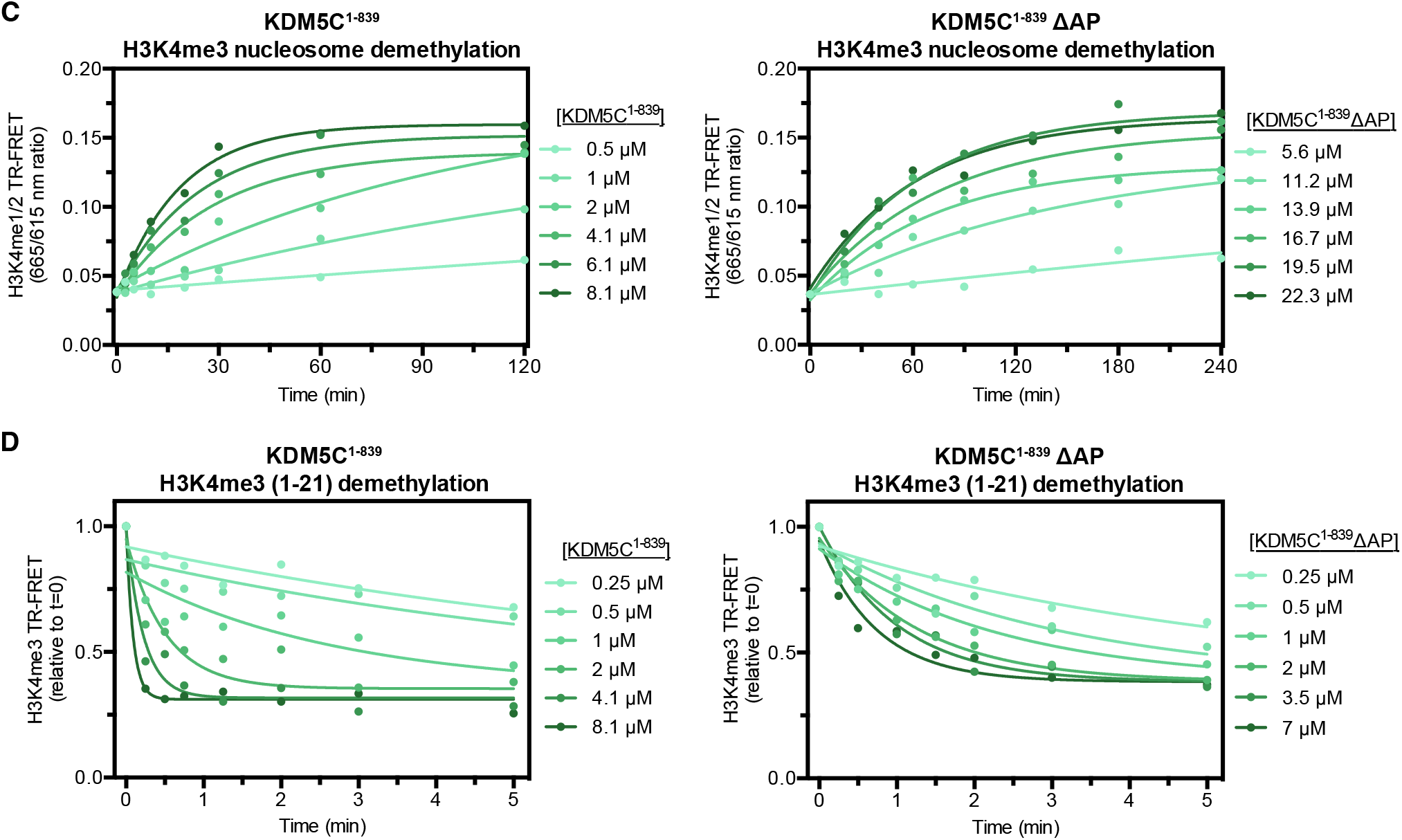
Related to Figure 1. **(A)** H3K4me3 substrate peptide demethylation by KDM5C constructs. *Left*: Demethylation kinetics of the H3K4me3 (1-21) substrate peptide by KDM5C constructs under single turnover conditions measured by a TR-FRET based kinetic assay. Observed rates are fit to a cooperative kinetic model, with *n* denoting the Hill coefficient. Representative kinetic traces used to determine observed demethylation rates are in Figure S1D. *Right*: Demethylation kinetics of the H3K4me3 (1-21) substrate peptide by KDM5C constructs under multiple turnover conditions measured by a formaldehyde release based kinetic assay. Deletion of the ARID and PHD1 region results in higher demethylase activity on the substrate peptide under multiple turnover conditions due to loss of substrate inhibition caused by this region. **(B)** Unmodified and substrate core nucleosome binding by KDM5C^1-839^ and KDM5C^1-839^ ΔAP. Nucleosome binding curves were measured by EMSA and fit to a cooperative binding model to determine apparent dissociation constants (*K*_d_^app^), with *n* denoting the Hill coefficient (*top*). Representative gel shifts of KDM5C binding to nucleosomes (*bottom*). Due to unattainable saturation of binding, a lower limit for the dissociation constant is presented for the unmodified nucleosome. **(C)** Representative demethylation kinetic traces of substrate nucleosome demethylation by KDM5C constructs (*left*: KDM5C^1-839^, *right*: KDM5C^1-839^ ΔAP) under single turnover conditions using TR-FRET based kinetic assay detecting formation of the H3K4me1/2 product nucleosome over time. Observed rates (*k*_obs_) are obtained by fitting kinetic traces to an exponential function. **(D)** Representative demethylation kinetic traces of substrate peptide demethylation by KDM5C constructs (*left*: KDM5C^1-839^, *right*: KDM5C^1-839^ ΔAP) under single turnover conditions using TR-FRET based kinetic assay detecting loss of the H3K4me3 substrate peptide over time. Observed rates (*k*_obs_) are obtained by fitting kinetic traces to an exponential function. All error bars represent SEM of at least two independent experiments (*n* ≥ 2).

**Figure S2.**
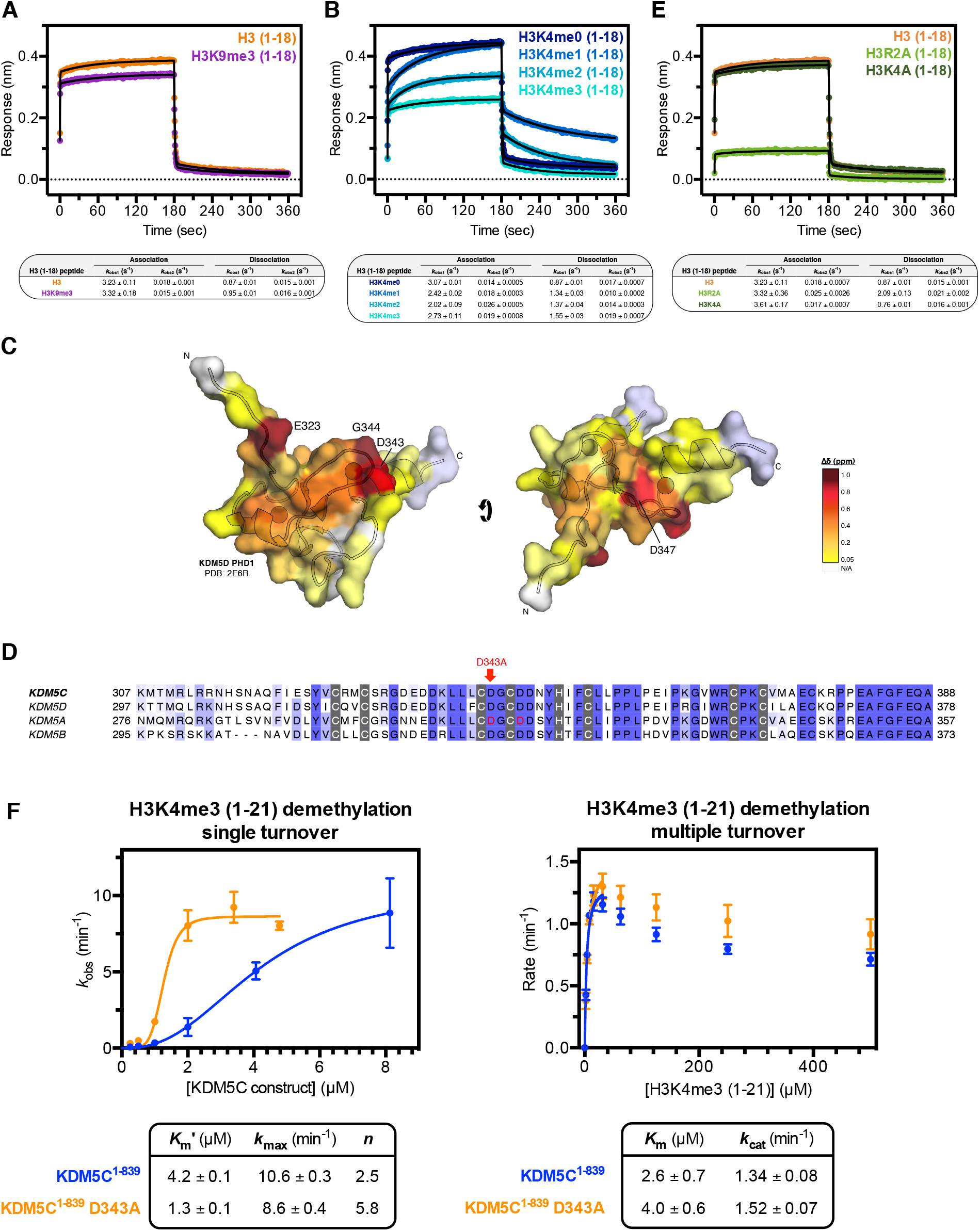
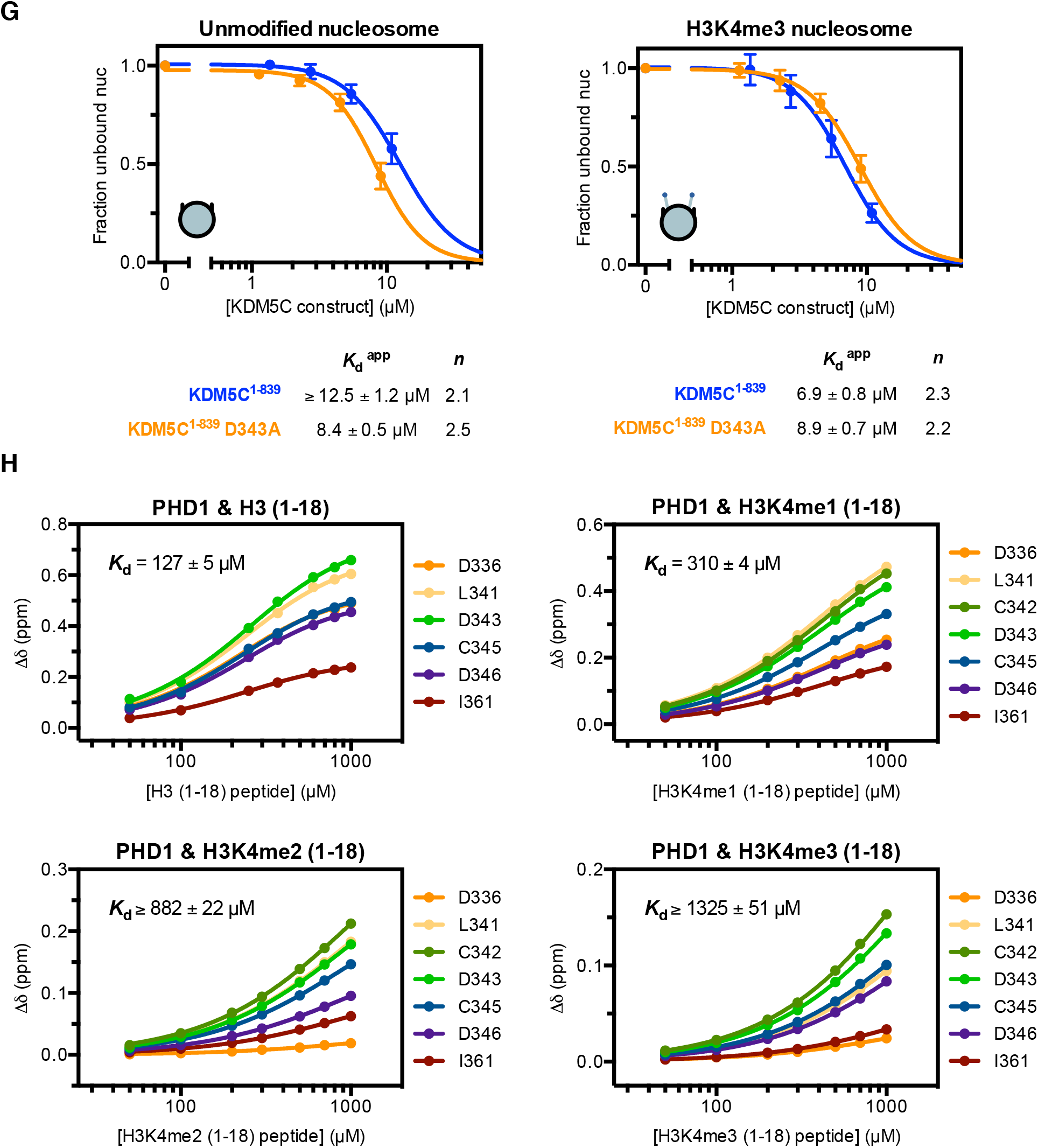
Related to Figure 2. **(A)** Binding kinetic trace of immobilized Avitag-PHD1 binding to H3 (1-18) and H3K9me3 (1-18) tail peptides measured by bio-layer interferometry (BLI). Observed rates (*k*_obs_) of association and dissociation are obtained by fitting kinetic traces to a two phase exponential function. **(B)** Binding kinetic trace of immobilized Avitag-PHD1 binding to H3K4me0/1/2/3 (1-18) tail peptides measured by BLI. Biphasic kinetic binding by PHD1 is modulated by the H3K4me state. **(C)** Chemical shift perturbations of PHD1 residues upon binding of the H3 (1-18) tail peptide (Figure 2A) colored by the gradient, unperturbed (yellow) to significantly perturbed (maroon), mapped to homologous residues in KDM5D PHD1 structure (PDB: 2E6R). Significantly perturbed residues are labeled. **(D)** Binding kinetic trace of immobilized Avitag-PHD1 binding to H3 (1-18) and H3 mutant (1-18) tail peptides (H3R2A and H3K4A) measured by BLI. Recognition of the H3 tail by PHD1 depends on the R2 residue but not K4 residue in H3. **(E)** Sequence alignment of PHD1 domains in KDM5A-D. The H3R2 recognizing residues D312 and D315 of KDM5A are indicated in red, and the PHD1 mutation D343A from this study is denoted above KDM5C. Zinc coordinating residues are highlighted in gray. **(F)** H3K4me3 substrate peptide demethylation by PHD1 mutant KDM5C^1-839^ relative to wild type. *Left*: Demethylation kinetics of the H3K4me3 (1-21) substrate peptide under single turnover conditions measured by a TR-FRET based kinetic assay. Observed rates are fit to a cooperative kinetic model, with *n* denoting the Hill coefficient. Unlike on the substrate nucleosome, the D343A PHD1 mutation does not increase catalytic rate on the substrate peptide, but does increase overall catalytic efficiency. *Right*: Demethylation kinetics of the H3K4me3 (1-21) substrate peptide under multiple turnover conditions measured by a formaldehyde release based kinetic assay. The D343A PHD1 mutation does not affect catalysis on the substrate peptide under these conditions, nor does it significantly affect substrate inhibition. **(G)** Unmodified and substrate core nucleosome binding by PHD1 mutant KDM5C^1-839^ relative to wild type. Nucleosome binding curves were measured by EMSA and fit to a cooperative binding model to determine apparent dissociation constants (*K*_d_^app^), with *n* denoting the Hill coefficient. Due to unattainable saturation of binding, a lower limit for the dissociation constant is presented for WT KDM5C binding the unmodified nucleosome. **(H)** Binding of the H3K4me0/1/2/3 (1-18) tail peptides by PHD1 by NMR titration HSQC experiments of indicated PHD1 residues that localize to the H3 binding surface (Figure S2C). Average dissociation constants with standard error for each ligand were determined from dissociation constants obtained from chemical shift changes (Δδ) of individual PHD1 residues. Due to incomplete saturation of binding, a lower limit for the dissociation constant is presented for the H3K4me2/3 peptides. All error bars represent SEM of at least two independent experiments (*n* ≥ 2).

**Figure S3.**
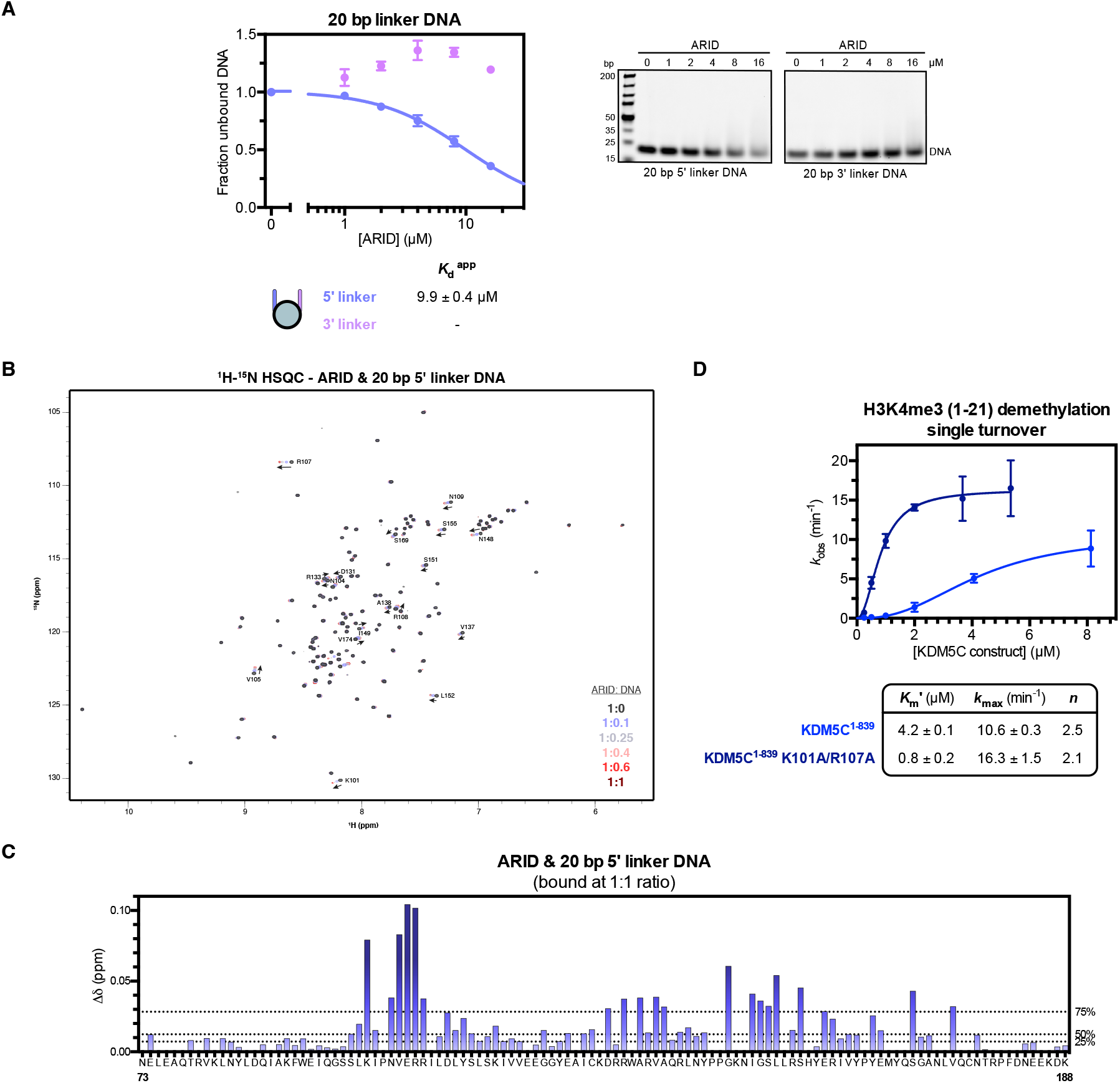
Related to Figure 3. **(A)** 20 bp linker DNA fragment binding by the ARID domain. Fragments contain 5’ and 3’ flanking DNA sequences used in the 187 bp nucleosome. Binding curves were measured by EMSA and fit to a binding model to determine apparent dissociation constants (*K*_*d*_ ^app^) (*left*). Representative gel shifts of ARID binding to 20 bp flanking linker DNA fragments (*right*). **(B)** 2D ^1^H-^15^N HSQC spectra of ARID titrated with increasing amounts of the 5’ linker DNA 20 bp fragment with indicated molar ratios. Assignments of most perturbed residues in ARID are labeled. **(C)** Chemical shift change (Δδ) of ARID residue backbone assignments upon binding of the 5’ linker DNA 20 bp fragment at 1:1 molar ratio measured by NMR. ARID backbone assignments could not be reliably transferred to a subset of residues and thus chemical shifts could not be determined (indicated by no values). Dashed lines indicate 25th, 50th, and 75th percentile rankings, and residues are colored by a gradient from unperturbed (light blue) to significantly perturbed (navy). **(D)** Demethylation kinetics of the H3K4me3 (1-21) substrate peptide by wild type and ARID mutant KDM5C^1-839^ under single turnover conditions. Observed rates are fit to a cooperative kinetic model, with *n* denoting the Hill coefficient. Unlike on the substrate nucleosome, the K101A/R107A ARID double mutation does not decrease catalytic rate on the substrate peptide, but does increase overall catalytic efficiency. All error bars represent SEM of at least two independent experiments (*n* ≥ 2).

**Figure S4.**
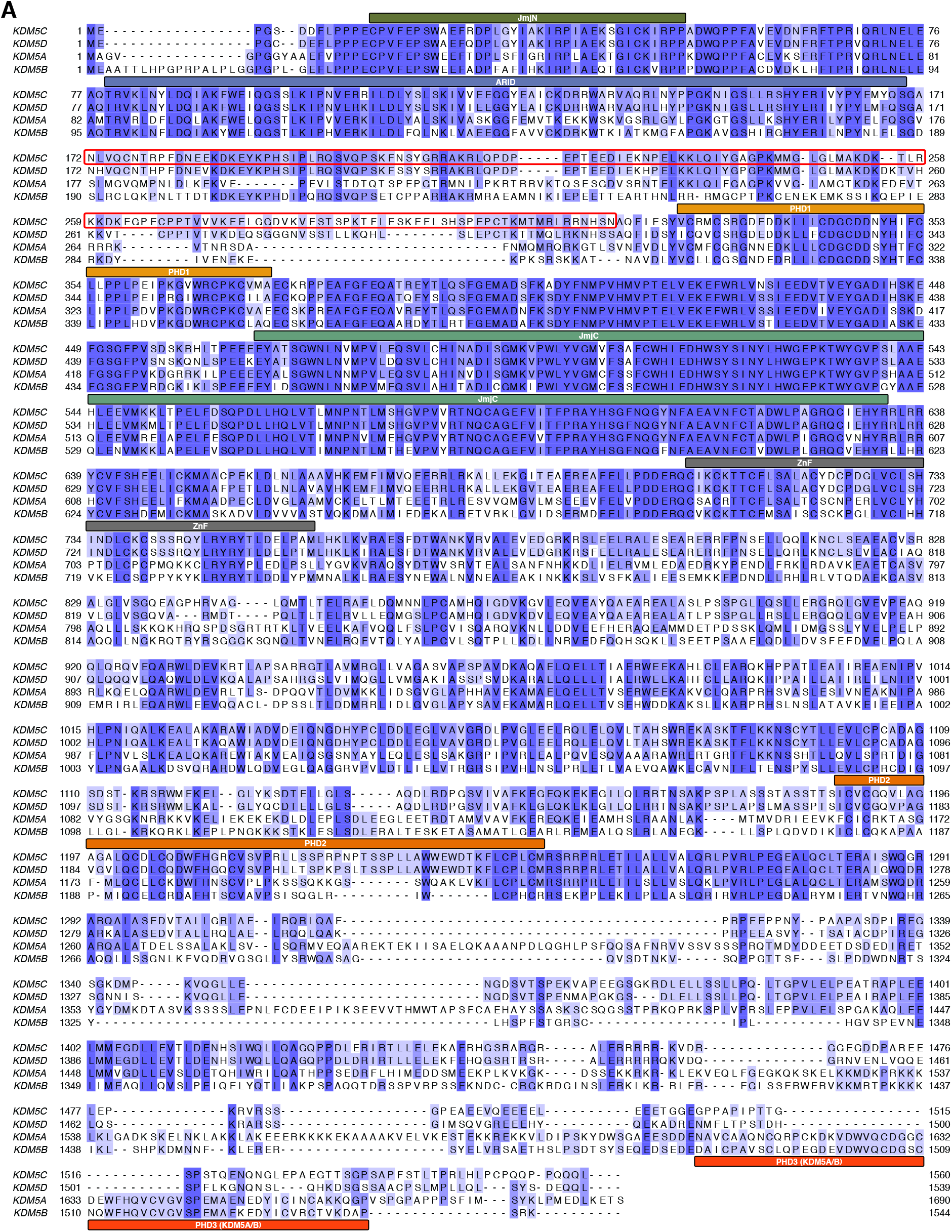
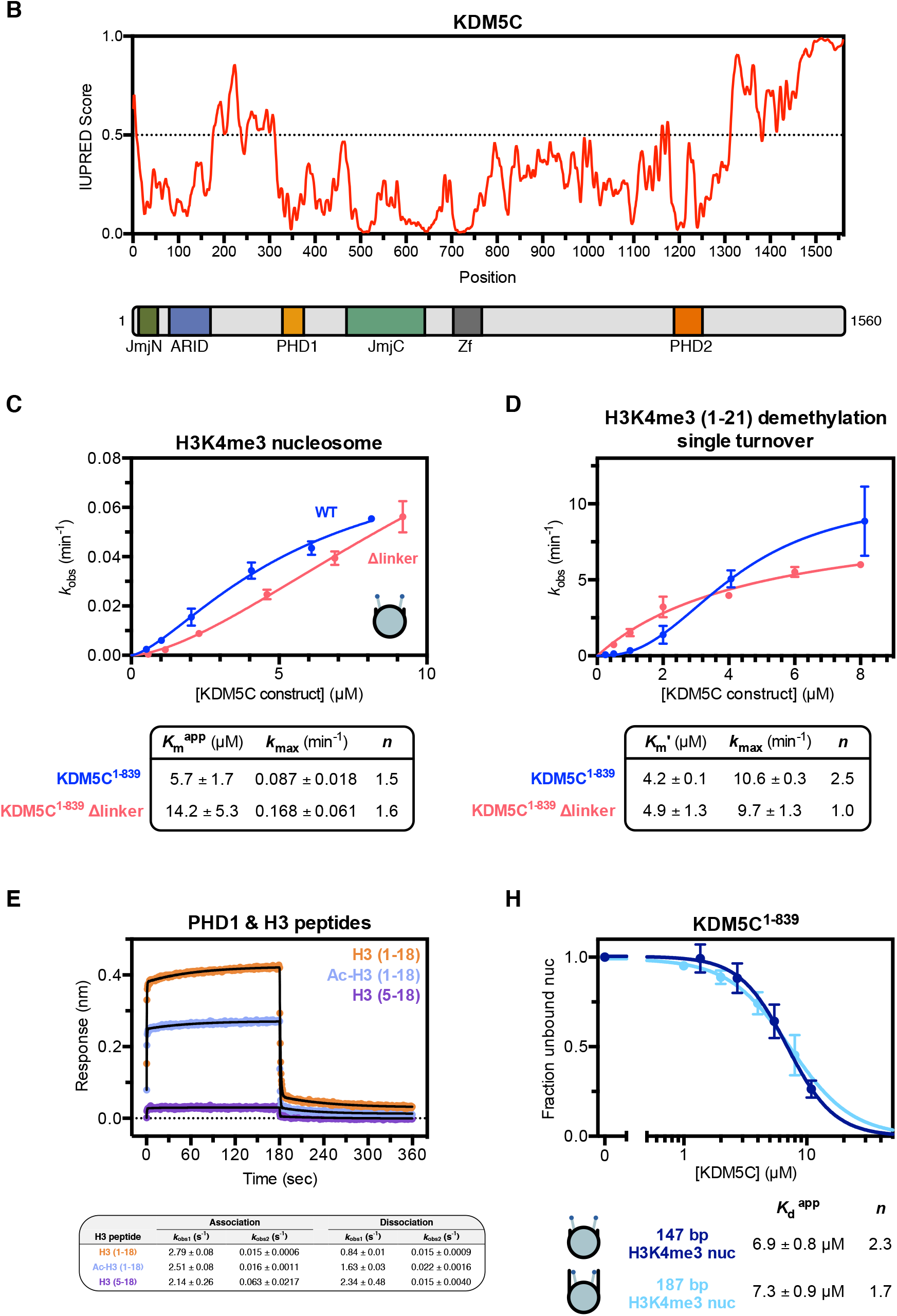
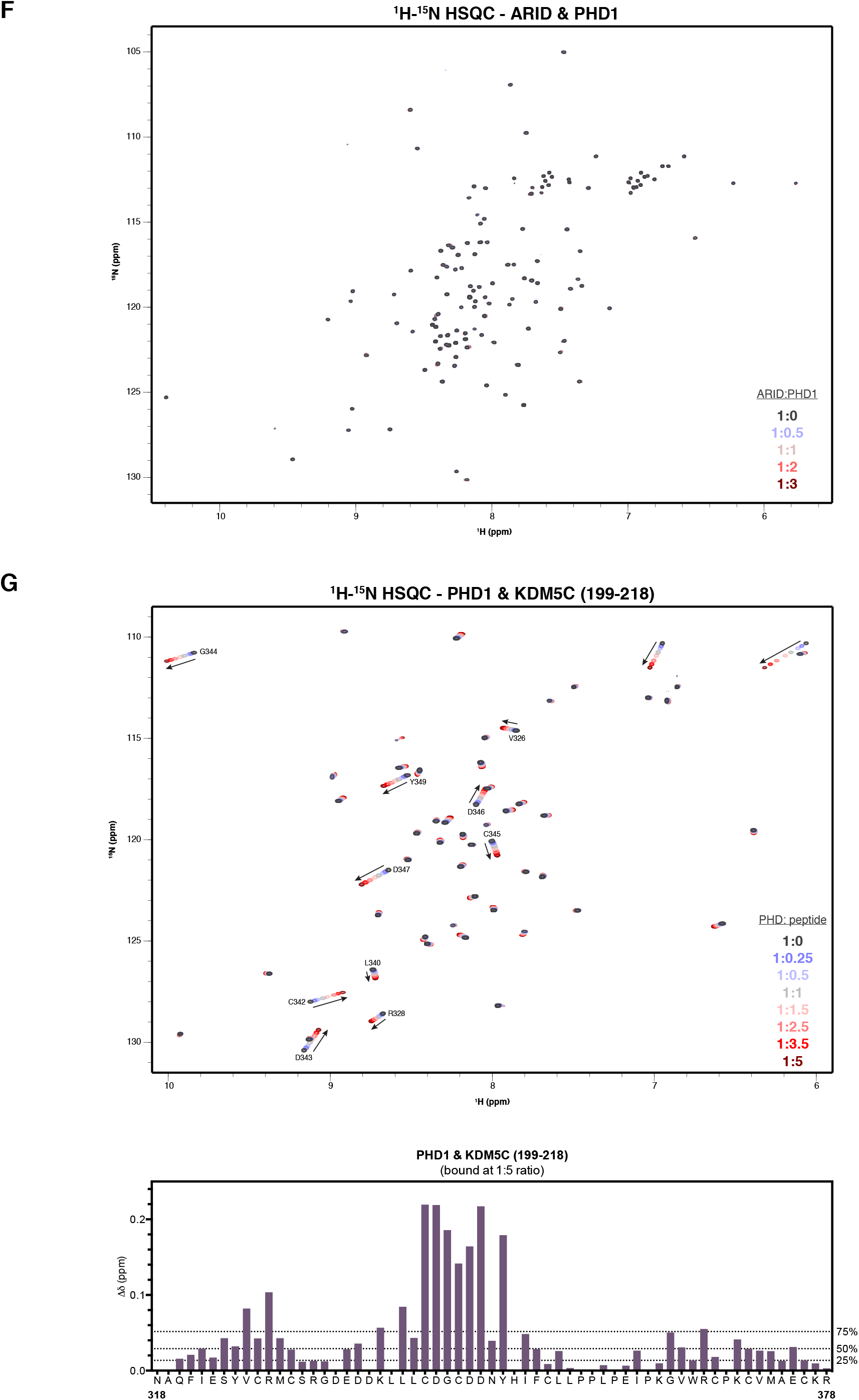
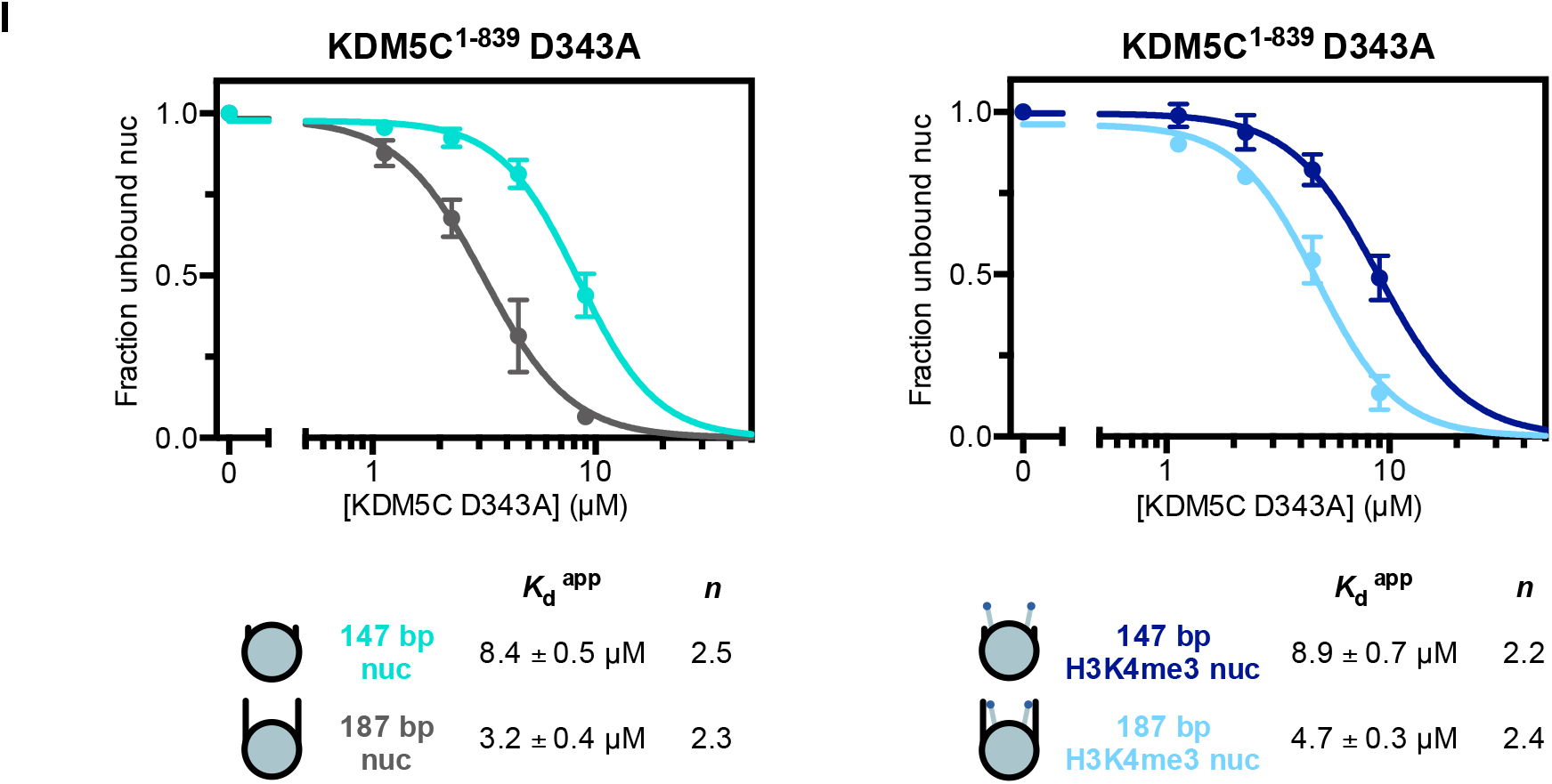
Related to Figure 4. **(A)** Sequence alignment of human KDM5A-D with annotated domains. KDM5C has a different and extended linker region between ARID and PHD1 (boxed in red). **(B)** IUPred profile [74] of predicted disorder in KDM5C (*top*) and annotated domain architecture of KDM5C (*bottom*). The linker between ARID and PHD1 is predicted to be disordered. **(C)** Demethylation kinetics of the H3K4me3 substrate nucleosome by wild type and KDM5C^1-839^ Δlinker under single turnover conditions. Observed rates are fit to a cooperative kinetic model, with *n* denoting the Hill coefficient. Deletion of the ARID-PHD1 linker does not significantly affect the catalytic efficiency of substrate nucleosome demethylation. **(D)** Demethylation kinetics of the H3K4me3 (1-21) substrate peptide by wild type and KDM5C^1-839^ Δlinker under single turnover conditions. Observed rates are fit to a cooperative kinetic model, with *n* denoting the Hill coefficient. Similarly to nucleosomes, deletion of the ARID-PHD1 linker does not significantly affect the catalytic efficiency of substrate peptide demethylation. **(E)** Binding kinetic trace of immobilized Avitag-PHD1 binding to H3 (1-18), N-terminally acetylated H3 (1-18), and H3 (5-18) tail peptides measured by bio-layer interferometry. Observed rates (*k*_obs_) of association and dissociation are obtained by fitting kinetic traces to a two phase exponential function. Recognition of the H3 tail by PHD1 does not strongly depend on the H3 N-terminus but does depend on the first 4 residues of H3 (ARTK). **(F)** 2D ^1^H-^15^N HSQC spectra of ARID titrated with increasing amounts of PHD1 with indicated molar ratios. PHD1 does not appear to bind the ARID domain. **(G)** 2D ^1^H-^15^N HSQC spectra of PHD1 titrated with increasing amounts of KDM5C (199-218) peptide with indicated molar ratios (*top*). Assignments of most perturbed residues in PHD1 are labeled. Corresponding chemical shift change (Δδ) of PHD1 residues upon binding of the KDM5C (199-218) peptide at 1:5 molar ratio (PHD:peptide) (*bottom*). Dashed lines indicate 25th, 50th, and 75th percentile rankings. **(H)** Nucleosome binding curves of KDM5C^1-839^ binding to substrate nucleosomes with and without 20 bp flanking DNA. Nucleosome binding curves were measured by EMSA and fit to a cooperative binding model to determine apparent dissociation constants (*K*_*d*_ ^app^), with *n* denoting the Hill coefficient. **(I)** Binding curves of PHD1 mutant KDM5C^1-839^ binding to unmodified and substrate nucleosomes with and without 20 bp flanking DNA. All error bars represent SEM of at least two independent experiments (*n* ≥ 2).

**Figure S5.**
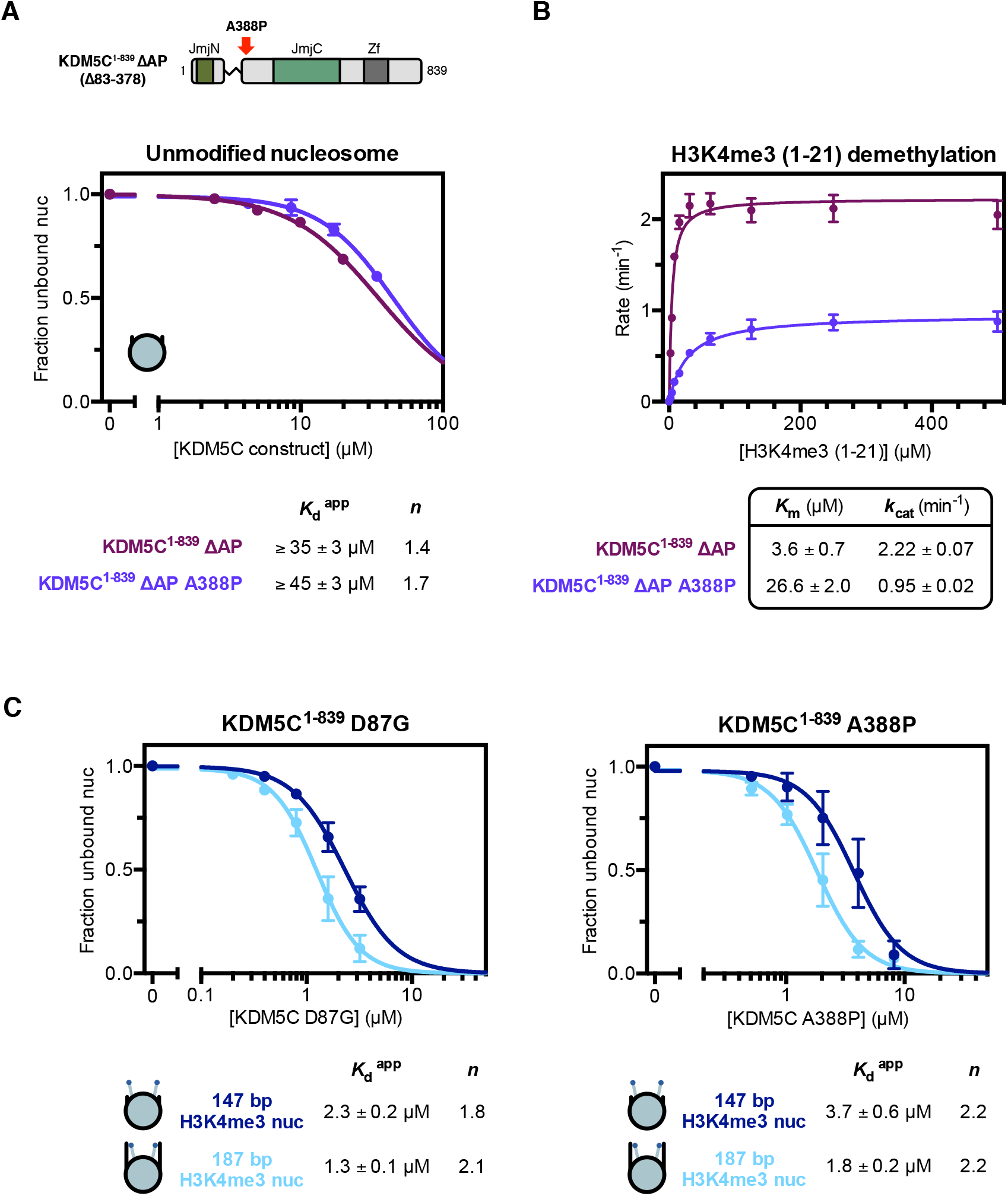
Related to Figure 5. **(A)** Unmodified core nucleosome binding by KDM5C^1-839^ ΔAP wild type and A388P. Nucleosome binding curves were measured by EMSA and fit to a cooperative binding model to determine apparent dissociation constants (*K*_d_^app^), with *n* denoting the Hill coefficient. Due to unattainable saturation of binding, a lower limit for the dissociation constant is presented. The A388P mutation does not enhance nucleosome binding in the absence of the ARID and PHD1 region, indicating this region in KDM5C is altered by the A388P mutation to enable enhanced binding. **(B)** Demethylation kinetics of the H3K4me3 (1-21) substrate peptide by KDM5C^1-839^ ΔAP wild type and A388P under multiple turnover conditions measured by a formaldehyde release based kinetic assay. The A388P mutation reduces demethylase activity of the catalytic domain alone, indicating distal structural disruption of the catalytic domain by this mutation. **(C)** Binding curves of KDM5C^1-839^ D87G and A388P binding to substrate nucleosomes with and without 20 bp flanking DNA. All error bars represent SEM of at least two independent experiments (*n* ≥ 2).

## REFERENCES

[1] H. Kimura, Histone modifications for human epigenome analysis, Journal of Human Genetics. 58 (2013) 439–445.

[2] S. Iwase, F. Lan, P. Bayliss, L. de la Torre-Ubieta, M. Huarte, H.H. Qi, J.R. Whetstine, A. Bonni, T.M. Roberts, Y. Shi, The X-linked mental retardation gene SMCX/JARID1C defines a family of histone H3 lysine 4 demethylases, Cell. 128 (2007) 1077–1088.

[3] R.J. Klose, Q. Yan, Z. Tothova, K. Yamane, H. Erdjument-Bromage, P. Tempst, D.G. Gilliland, Y. Zhang, W.G. Kaelin Jr, The retinoblastoma binding protein RBP2 is an H3K4 demethylase, Cell. 128 (2007) 889–900.

[4] K. Yamane, K. Tateishi, R.J. Klose, J. Fang, L.A. Fabrizio, H. Erdjument-Bromage, J. Taylor-Papadimitriou, P. Tempst, Y. Zhang, PLU-1 is an H3K4 demethylase involved in transcriptional repression and breast cancer cell proliferation, Mol. Cell. 25 (2007) 801–812.

[5] M.G. Lee, J. Norman, A. Shilatifard, R. Shiekhattar, Physical and functional association of a trimethyl H3K4 demethylase and Ring6a/MBLR, a polycomb-like protein, Cell. 128 (2007) 877–887.

[6] Y. Shi, F. Lan, C. Matson, P. Mulligan, J.R. Whetstine, P.A. Cole, R.A. Casero, Y. Shi, Histone demethylation mediated by the nuclear amine oxidase homolog LSD1, Cell. 119 (2004) 941–953.

[7] J. Christensen, K. Agger, P.A.C. Cloos, D. Pasini, S. Rose, L. Sennels, J. Rappsilber, K.H. Hansen, A.E. Salcini, K. Helin, RBP2 belongs to a family of demethylases, specific for tri- and dimethylated lysine 4 on histone 3, Cell. 128 (2007) 1063–1076.

[8] J.R. Horton, A. Engstrom, E.L. Zoeller, X. Liu, J.R. Shanks, X. Zhang, M.A. Johns, P.M. Vertino, H. Fu, X. Cheng, Characterization of a Linked Jumonji Domain of the KDM5/JARID1 Family of Histone H3 Lysine 4 Demethylases, J. Biol. Chem. 291 (2016) 2631–2646.

[9] C. Johansson, S. Velupillai, A. Tumber, A. Szykowska, E.S. Hookway, R.P. Nowak, C. Strain-Damerell, C. Gileadi, M. Philpott, N. Burgess-Brown, N. Wu, J. Kopec, A. Nuzzi, H. Steuber, U. Egner, V. Badock, S. Munro, N.B. LaThangue, S. Westaway, J. Brown, N. Athanasou, R. Prinjha, P.E. Brennan, U. Oppermann, Structural analysis of human KDM5B guides histone demethylase inhibitor development, Nat. Chem. Biol. 12 (2016) 539–545.

[10] L. Li, C. Greer, R.N. Eisenman, J. Secombe, Essential functions of the histone demethylase lid, PLoS Genet. 6 (2010) e1001221.

[11] E. Brookes, B. Laurent, K. Õunap, R. Carroll, J.B. Moeschler, M. Field, C.E. Schwartz, J. Gecz, Y. Shi, Mutations in the intellectual disability gene KDM5C reduce protein stability and demethylase activity, Hum. Mol. Genet. 24 (2015) 2861–2872.

[12] Y. Xiang, Z. Zhu, G. Han, X. Ye, B. Xu, Z. Peng, Y. Ma, Y. Yu, H. Lin, A.P. Chen, C.D. Chen, JARID1B is a histone H3 lysine 4 demethylase up-regulated in prostate cancer, Proc. Natl. Acad. Sci. U. S. A. 104 (2007) 19226–19231.

[13] S. Tu, Y.-C. Teng, C. Yuan, Y.-T. Wu, M.-Y. Chan, A.-N. Cheng, P.-H. Lin, L.-J. Juan, M.-D. Tsai, The ARID domain of the H3K4 demethylase RBP2 binds to a DNA CCGCCC motif, Nat. Struct. Mol. Biol. 15 (2008) 419–421.

[14] A.G. Scibetta, S. Santangelo, J. Coleman, D. Hall, T. Chaplin, J. Copier, S. Catchpole, J. Burchell, J. Taylor-Papadimitriou, Functional analysis of the transcription repressor PLU-1/JARID1B, Mol. Cell. Biol. 27 (2007) 7220–7235.

[15] W. Yao, Y. Peng, D. Lin, The flexible loop L1 of the H3K4 demethylase JARID1B ARID domain has a crucial role in DNA-binding activity, Biochem. Biophys. Res. Commun. 396 (2010) 323–328.

[16] F. Lan, R.E. Collins, R. De Cegli, R. Alpatov, J.R. Horton, X. Shi, O. Gozani, X. Cheng, Y. Shi, Recognition of unmethylated histone H3 lysine 4 links BHC80 to LSD1-mediated gene repression, Nature. 448 (2007) 718–722.

[17] B.J. Klein, X. Wang, G. Cui, C. Yuan, M.V. Botuyan, K. Lin, Y. Lu, X. Wang, Y. Zhao, C.J. Bruns, G. Mer, X. Shi, T.G. Kutateladze, PHF20 Readers Link Methylation of Histone H3K4 and p53 with H4K16 Acetylation, Cell Rep. 17 (2016) 1158–1170.

[18] X. Shi, T. Hong, K.L. Walter, M. Ewalt, E. Michishita, T. Hung, D. Carney, P. Peña, F. Lan, M.R. Kaadige, N. Lacoste, C. Cayrou, F. Davrazou, A. Saha, B.R. Cairns, D.E. Ayer, T.G. Kutateladze, Y. Shi, J. Côté, K.F. Chua, O. Gozani, ING2 PHD domain links histone H3 lysine 4 methylation to active gene repression, Nature. 442 (2006) 96–99.

[19] H. Li, S. Ilin, W. Wang, E.M. Duncan, J. Wysocka, C.D. Allis, D.J. Patel, Molecular basis for site-specific read-out of histone H3K4me3 by the BPTF PHD finger of NURF, Nature. 442 (2006) 91–95.

[20] P.V. Peña, F. Davrazou, X. Shi, K.L. Walter, V.V. Verkhusha, O. Gozani, R. Zhao, T.G. Kutateladze, Molecular mechanism of histone H3K4me3 recognition by plant homeodomain of ING2, Nature. 442 (2006) 100–103.

[21] J. Wysocka, T. Swigut, H. Xiao, T.A. Milne, S.Y. Kwon, J. Landry, M. Kauer, A.J. Tackett, B.T. Chait, P. Badenhorst, C. Wu, C.D. Allis, A PHD finger of NURF couples histone H3 lysine 4 trimethylation with chromatin remodelling, Nature. 442 (2006) 86–90.

[22] R. Sanchez, M.-M. Zhou, The PHD finger: a versatile epigenome reader, Trends Biochem. Sci. 36 (2011) 364–372.

[23] B.J. Klein, L. Piao, Y. Xi, H. Rincon-Arano, S.B. Rothbart, D. Peng, H. Wen, C. Larson, X. Zhang, X. Zheng, M.A. Cortazar, P.V. Peña, A. Mangan, D.L. Bentley, B.D. Strahl, M. Groudine, W. Li, X. Shi, T.G. Kutateladze, The histone-H3K4-specific demethylase KDM5B binds to its substrate and product through distinct PHD fingers, Cell Rep. 6 (2014) 325– 335.

[24] Y. Zhang, H. Yang, X. Guo, N. Rong, Y. Song, Y. Xu, W. Lan, X. Zhang, M. Liu, Y. Xu, C. Cao, The PHD1 finger of KDM5B recognizes unmodified H3K4 during the demethylation of histone H3K4me2/3 by KDM5B, Protein Cell. 5 (2014) 837–850.

[25] I.O. Torres, K.M. Kuchenbecker, C.I. Nnadi, R.J. Fletterick, M.J.S. Kelly, D.G. Fujimori, Histone demethylase KDM5A is regulated by its reader domain through a positive-feedback mechanism, Nat. Commun. 6 (2015) 6204.

[26] S. Zhao, K.N. Chuh, B. Zhang, B.E. Dul, R.E. Thompson, L.A. Farrelly, X. Liu, N. Xu, Y. Xue, R.G. Roeder, I. Maze, T.W. Muir, H. Li, Histone H3Q5 serotonylation stabilizes H3K4 methylation and potentiates its readout, Proc. Natl. Acad. Sci. U. S. A. 118 (2021). https://doi.org/10.1073/pnas.2016742118.

[27] G.G. Wang, J. Song, Z. Wang, H.L. Dormann, F. Casadio, H. Li, J.-L. Luo, D.J. Patel, C.D. Allis, Haematopoietic malignancies caused by dysregulation of a chromatin-binding PHD finger, Nature. 459 (2009) 847–851.

[28] J.E. Longbotham, C.M. Chio, V. Dharmarajan, M.J. Trnka, I.O. Torres, D. Goswami, K. Ruiz, A.L. Burlingame, P.R. Griffin, D.G. Fujimori, Histone H3 binding to the PHD1 domain of histone demethylase KDM5A enables active site remodeling, Nat. Commun. 10 (2019) 94.

[29] S. Iwase, E. Brookes, S. Agarwal, A.I. Badeaux, H. Ito, C.N. Vallianatos, G.S. Tomassy, T. Kasza, G. Lin, A. Thompson, L. Gu, K.Y. Kwan, C. Chen, M.A. Sartor, B. Egan, J. Xu, Y. Shi, A Mouse Model of X-linked Intellectual Disability Associated with Impaired Removal of Histone Methylation, Cell Rep. 14 (2016) 1000–1009.

[30] L.R. Jensen, H. Bartenschlager, S. Rujirabanjerd, A. Tzschach, A. Nümann, A.R. Janecke, R. Spörle, S. Stricker, M. Raynaud, J. Nelson, A. Hackett, J.-P. Fryns, J. Chelly, A.P. de Brouwer, B. Hamel, J. Gecz, H.-H. Ropers, A.W. Kuss, A distinctive gene expression fingerprint in mentally retarded male patients reflects disease-causing defects in the histone demethylase KDM5C, Pathogenetics. 3 (2010) 2.

[31] L.R. Jensen, M. Amende, U. Gurok, B. Moser, V. Gimmel, A. Tzschach, A.R. Janecke, G. Tariverdian, J. Chelly, J.-P. Fryns, H. Van Esch, T. Kleefstra, B. Hamel, C. Moraine, J. Gecz, G. Turner, R. Reinhardt, V.M. Kalscheuer, H.-H. Ropers, S. Lenzner, Mutations in the JARID1C gene, which is involved in transcriptional regulation and chromatin remodeling, cause X-linked mental retardation, Am. J. Hum. Genet. 76 (2005) 227–236.

[32] M. Scandaglia, J.P. Lopez-Atalaya, A. Medrano-Fernandez, M.T. Lopez-Cascales, B. Del Blanco, M. Lipinski, E. Benito, R. Olivares, S. Iwase, Y. Shi, A. Barco, Loss of Kdm5c Causes Spurious Transcription and Prevents the Fine-Tuning of Activity-Regulated Enhancers in Neurons, Cell Rep. 21 (2017) 47–59.

[33] N.S. Outchkourov, J.M. Muiño, K. Kaufmann, W.F.J. van Ijcken, M.J. Groot Koerkamp, D. van Leenen, P. de Graaf, F.C.P. Holstege, F.G. Grosveld, H.T.M. Timmers, Balancing of histone H3K4 methylation states by the Kdm5c/SMCX histone demethylase modulates promoter and enhancer function, Cell Rep. 3 (2013) 1071–1079.

[34] H. Shen, W. Xu, R. Guo, B. Rong, L. Gu, Z. Wang, C. He, L. Zheng, X. Hu, Z. Hu, Z.-M. Shao, P. Yang, F. Wu, Y.G. Shi, Y. Shi, F. Lan, Suppression of Enhancer Overactivation by a RACK7-Histone Demethylase Complex, Cell. 165 (2016) 331–342.

[35] T.F. Gonçalves, A.P. Gonçalves, N. Fintelman Rodrigues, J.M. dos Santos, M.M.G. Pimentel, C.B. Santos-Rebouças, KDM5C mutational screening among males with intellectual disability suggestive of X-Linked inheritance and review of the literature, Eur. J. Med. Genet. 57 (2014) 138–144.

[36] A. Tzschach, S. Lenzner, B. Moser, R. Reinhardt, J. Chelly, J.-P. Fryns, T. Kleefstra, M. Raynaud, G. Turner, H.-H. Ropers, A. Kuss, L.R. Jensen, Novel JARID1C/SMCX mutations in patients with X-linked mental retardation, Hum. Mutat. 27 (2006) 389.

[37] C. Santos, L. Rodriguez-Revenga, I. Madrigal, C. Badenas, M. Pineda, M. Milà, A novel mutation in JARID1C gene associated with mental retardation, Eur. J. Hum. Genet. 14 (2006) 583–586.

[38] F.E. Abidi, L. Holloway, C.A. Moore, D.D. Weaver, R.J. Simensen, R.E. Stevenson, R.C. Rogers, C.E. Schwartz, Mutations in JARID1C are associated with X-linked mental retardation, short stature and hyperreflexia, J. Med. Genet. 45 (2008) 787–793.

[39] S. Rujirabanjerd, J. Nelson, P.S. Tarpey, A. Hackett, S. Edkins, F. Lucy Raymond, C.E. Schwartz, G. Turner, S. Iwase, Y. Shi, P. Andrew Futreal, M.R. Stratton, J. Gecz, Identification and characterization of two novel JARID1C mutations: suggestion of an emerging genotype–phenotype correlation, European Journal of Human Genetics. 18 (2010) 330–335. https://doi.org/10.1038/ejhg.2009.175.

[40] J. Xu, P.S. Burgoyne, A.P. Arnold, Sex differences in sex chromosome gene expression in mouse brain, Human Molecular Genetics. 11 (2002) 1409–1419. https://doi.org/10.1093/hmg/11.12.1409.

[41] C.N. Vallianatos, C. Farrehi, M.J. Friez, M. Burmeister, C.E. Keegan, S. Iwase, Altered Gene-Regulatory Function of KDM5C by a Novel Mutation Associated With Autism and Intellectual Disability, Front. Mol. Neurosci. 11 (2018) 104.

[42] M. Tahiliani, P. Mei, R. Fang, T. Leonor, M. Rutenberg, F. Shimizu, J. Li, A. Rao, Y. Shi, The histone H3K4 demethylase SMCX links REST target genes to X-linked mental retardation, Nature. 447 (2007) 601–605.

[43] B.N. Jones, D.-U. Quang-Dang, Y. Oku, J.D. Gross, A kinetic assay to monitor RNA decapping under single-turnover conditions, Methods Enzymol. 448 (2008) 23–40.

[44] J.E. Longbotham, M.J.S. Kelly, D.G. Fujimori, Recognition of Histone H3 Methylation States by the PHD1 Domain of Histone Demethylase KDM5A, ACS Chem. Biol. (2021). https://doi.org/10.1021/acschembio.0c00976.

[45] C. Koehler, S. Bishop, E.F. Dowler, P. Schmieder, A. Diehl, H. Oschkinat, L.J. Ball, Backbone and sidechain 1H, 13C and 15N resonance assignments of the Bright/ARID domain from the human JARID1C (SMCX) protein, Biomol. NMR Assign. 2 (2008) 9–11.

[46] L. Poeta, A. Padula, M.B. Lioi, H. van Bokhoven, M.G. Miano, Analysis of a Set of KDM5C Regulatory Genes Mutated in Neurodevelopmental Disorders Identifies Temporal Coexpression Brain Signatures, Genes. 12 (2021). https://doi.org/10.3390/genes12071088.

[47] S.-A. Kim, J. Zhu, N. Yennawar, P. Eek, S. Tan, Crystal Structure of the LSD1/CoREST Histone Demethylase Bound to Its Nucleosome Substrate, Mol. Cell. 78 (2020) 903–914.e4.

[48] V. Kasinath, C. Beck, P. Sauer, S. Poepsel, J. Kosmatka, M. Faini, D. Toso, R. Aebersold, E. Nogales, JARID2 and AEBP2 regulate PRC2 in the presence of H2AK119ub1 and other histone modifications, Science. 371 (2021). https://doi.org/10.1126/science.abc3393.

[49] K. Finogenova, J. Bonnet, S. Poepsel, I.B. Schäfer, K. Finkl, K. Schmid, C. Litz, M. Strauss, C. Benda, J. Müller, Structural basis for PRC2 decoding of active histone methylation marks H3K36me2/3, Elife. 9 (2020). https://doi.org/10.7554/eLife.61964.

[50] E.J. Worden, X. Zhang, C. Wolberger, Structural basis for COMPASS recognition of an H2B-ubiquitinated nucleosome, Elife. 9 (2020). https://doi.org/10.7554/eLife.53199.

[51] W. Li, W. Tian, G. Yuan, P. Deng, D. Sengupta, Z. Cheng, Y. Cao, J. Ren, Y. Qin, Y. Zhou, Y. Jia, O. Gozani, D.J. Patel, Z. Wang, Molecular basis of nucleosomal H3K36 methylation by NSD methyltransferases, Nature. 590 (2021) 498–503.

[52] S. Bilokapic, M.J. Suskiewicz, I. Ahel, M. Halic, Bridging of DNA breaks activates PARP2– HPF1 to modify chromatin, Nature. 585 (2020) 609–613. https://doi.org/10.1038/s41586-020-2725-7.

[53] S. Bilokapic, M. Halic, Nucleosome and ubiquitin position Set2 to methylate H3K36, Nat. Commun. 10 (2019) 3795.

[54] P.L. Hsu, H. Shi, C. Leonen, J. Kang, C. Chatterjee, N. Zheng, Structural Basis of H2B Ubiquitination-Dependent H3K4 Methylation by COMPASS, Mol. Cell. 76 (2019) 712–723.e4.

[55] C. Marabelli, B. Marrocco, S. Pilotto, S. Chittori, S. Picaud, S. Marchese, G. Ciossani, F. Forneris, P. Filippakopoulos, G. Schoehn, D. Rhodes, S. Subramaniam, A. Mattevi, A Tail-Based Mechanism Drives Nucleosome Demethylation by the LSD2/NPAC Multimeric Complex, Cell Rep. 27 (2019) 387–399.e7.

[56] S. Poepsel, V. Kasinath, E. Nogales, Cryo-EM structures of PRC2 simultaneously engaged with two functionally distinct nucleosomes, Nat. Struct. Mol. Biol. 25 (2018) 154–162.

[57] S.H. Park, A. Ayoub, Y.-T. Lee, J. Xu, H. Kim, W. Zheng, B. Zhang, L. Sha, S. An, Y. Zhang, M.A. Cianfrocco, M. Su, Y. Dou, U.-S. Cho, Cryo-EM structure of the human MLL1 core complex bound to the nucleosome, Nat. Commun. 10 (2019) 1–13.

[58] H. Xue, T. Yao, M. Cao, G. Zhu, Y. Li, G. Yuan, Y. Chen, M. Lei, J. Huang, Structural basis of nucleosome recognition and modification by MLL methyltransferases, Nature. 573 (2019) 445–449.

[59] S. Hatazawa, J. Liu, Y. Takizawa, M. Zandian, L. Negishi, T.G. Kutateladze, H. Kurumizaka, Structural basis for binding diversity of acetyltransferase p300 to the nucleosome, iScience. 25 (2022) 104563. https://doi.org/10.1016/j.isci.2022.104563.

[60] J. Gatchalian, X. Wang, J. Ikebe, K.L. Cox, A.H. Tencer, Y. Zhang, N.L. Burge, L. Di, M.D. Gibson, C.A. Musselman, M.G. Poirier, H. Kono, J.J. Hayes, T.G. Kutateladze, Accessibility of the histone H3 tail in the nucleosome for binding of paired readers, Nat. Commun. 8 (2017) 1489.

[61] E.A. Morrison, S. Bowerman, K.L. Sylvers, J. Wereszczynski, C.A. Musselman, The conformation of the histone H3 tail inhibits association of the BPTF PHD finger with the nucleosome, Elife. 7 (2018). https://doi.org/10.7554/eLife.31481.

[62] A. Stützer, S. Liokatis, A. Kiesel, D. Schwarzer, R. Sprangers, J. Söding, P. Selenko, W. Fischle, Modulations of DNA Contacts by Linker Histones and Post-translational Modifications Determine the Mobility and Modifiability of Nucleosomal H3 Tails, Mol. Cell. 61 (2016) 247–259.

[63] Y. Peng, S. Li, A. Onufriev, D. Landsman, A.R. Panchenko, Binding of regulatory proteins to nucleosomes is modulated by dynamic histone tails, Nat. Commun. 12 (2021) 5280.

[64] T.M. Weaver, E.A. Morrison, C.A. Musselman, Reading More than Histones: The Prevalence of Nucleic Acid Binding among Reader Domains, Molecules. 23 (2018). https://doi.org/10.3390/molecules23102614.

[65] S. Pilotto, V. Speranzini, M. Tortorici, D. Durand, A. Fish, S. Valente, F. Forneris, A. Mai, T.K. Sixma, P. Vachette, A. Mattevi, Interplay among nucleosomal DNA, histone tails, and corepressor CoREST underlies LSD1-mediated H3 demethylation, Proc. Natl. Acad. Sci. U. S. A. 112 (2015) 2752–2757.

[66] C.A. Musselman, T.G. Kutateladze, Characterization of functional disordered regions within chromatin-associated proteins, iScience. 24 (2021) 102070.

[67] S. Contreras-Martos, A. Piai, S. Kosol, M. Varadi, A. Bekesi, P. Lebrun, A.N. Volkov, K. Gevaert, R. Pierattelli, I.C. Felli, P. Tompa, Linking functions: an additional role for an intrinsically disordered linker domain in the transcriptional coactivator CBP, Sci. Rep. 7 (2017) 4676.

[68] L. Jiao, M. Shubbar, X. Yang, Q. Zhang, S. Chen, Q. Wu, Z. Chen, J. Rizo, X. Liu, A partially disordered region connects gene repression and activation functions of EZH2, Proc. Natl. Acad. Sci. U. S. A. 117 (2020) 16992–17002.

[69] M.M. Keenen, D. Brown, L.D. Brennan, R. Renger, H. Khoo, C.R. Carlson, B. Huang, S.W. Grill, G.J. Narlikar, S. Redding, HP1 proteins compact DNA into mechanically and positionally stable phase separated domains, Elife. 10 (2021). https://doi.org/10.7554/eLife.64563.

[70] S.-A. Kim, N. Chatterjee, M.J. Jennings, B. Bartholomew, S. Tan, Extranucleosomal DNA enhances the activity of the LSD1/CoREST histone demethylase complex, Nucleic Acids Res. 43 (2015) 4868–4880.

[71] J.C. Zhou, N.P. Blackledge, A.M. Farcas, R.J. Klose, Recognition of CpG island chromatin by KDM2A requires direct and specific interaction with linker DNA, Mol. Cell. Biol. 32 (2012) 479–489.

[72] N.P. Blackledge, J.C. Zhou, M.Y. Tolstorukov, A.M. Farcas, P.J. Park, R.J. Klose, CpG islands recruit a histone H3 lysine 36 demethylase, Mol. Cell. 38 (2010) 179–190.

[73] S. Zamurrad, H.A.M. Hatch, C. Drelon, H.M. Belalcazar, J. Secombe, A Drosophila Model of Intellectual Disability Caused by Mutations in the Histone Demethylase KDM5, Cell Rep. 22 (2018) 2359–2369.

[74] G. Erdős, M. Pajkos, Z. Dosztányi, IUPred3: prediction of protein disorder enhanced with unambiguous experimental annotation and visualization of evolutionary conservation, Nucleic Acids Res. 49 (2021) W297–W303.

