## Supplemental Information for "Chromatin sensing by the auxiliary domains of KDM5C regulates its demethylase activity and is disrupted by X-linked intellectual disability mutations"

Figure S1

Figure S2

Figure S3

Figure S4

Figure S5

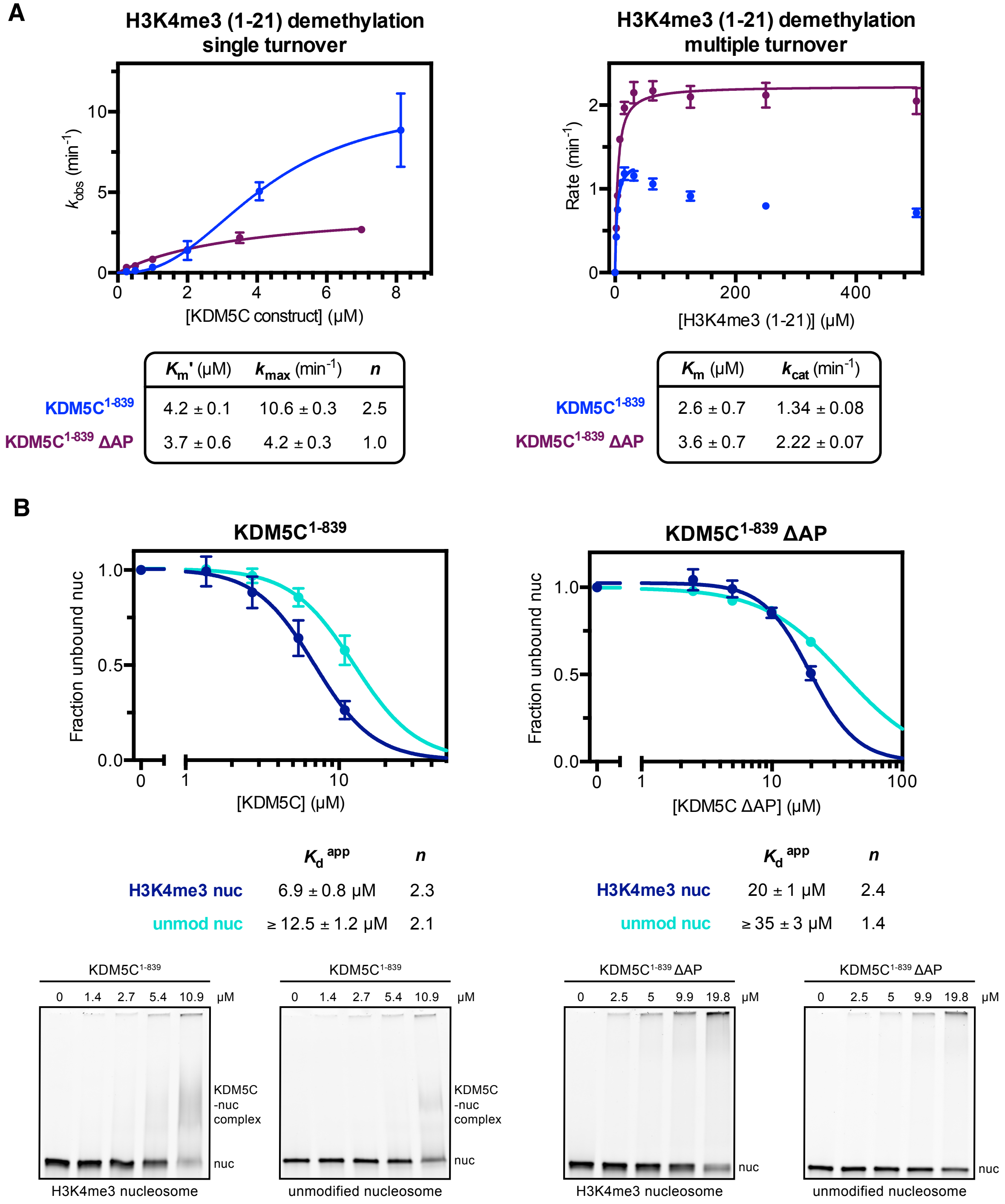

A
H3K4me3 (1-21) demethylation single turnover
k_obs (min-1)
[KDM5C construct] (μM)
H3K4me3 (1-21) demethylation multiple turnover
Rate (min-1)
[H3K4me3 (1-21)] (μM)
Km' (μM) kmax (min-1) n
KDM5C1-839 4.2 ± 0.1 10.6 ± 0.3 2.5
KDM5C1-839 ΔAP 3.7 ± 0.6 4.2 ± 0.3 1.0
Km (μM) kcat (min-1)
KDM5C1-839 2.6 ± 0.7 1.34 ± 0.08
KDM5C1-839 ΔAP 3.6 ± 0.7 2.22 ± 0.07
B
KDM5C1-839
Fraction unbound nuc
[KDM5C] (μM)
KDM5C1-839 ΔAP
[KDM5C ΔAP] (μM)
Kd app n
H3K4me3 nuc 6.9 ± 0.8 μM 2.3
unmod nuc ≥ 12.5 ± 1.2 μM 2.1
Kd app n
H3K4me3 nuc 20 ± 1 μM 2.4
unmod nuc ≥ 35 ± 3 μM 1.4
KDM5C1-839
0 1.4 2.7 5.4 10.9 μM
KDM5C -nuc complex
nuc
H3K4me3 nucleosome
unmodified nucleosome
KDM5C1-839 ΔAP
0 2.5 5 9.9 19.8 μM
H3K4me3 nucleosome
unmodified nucleosome

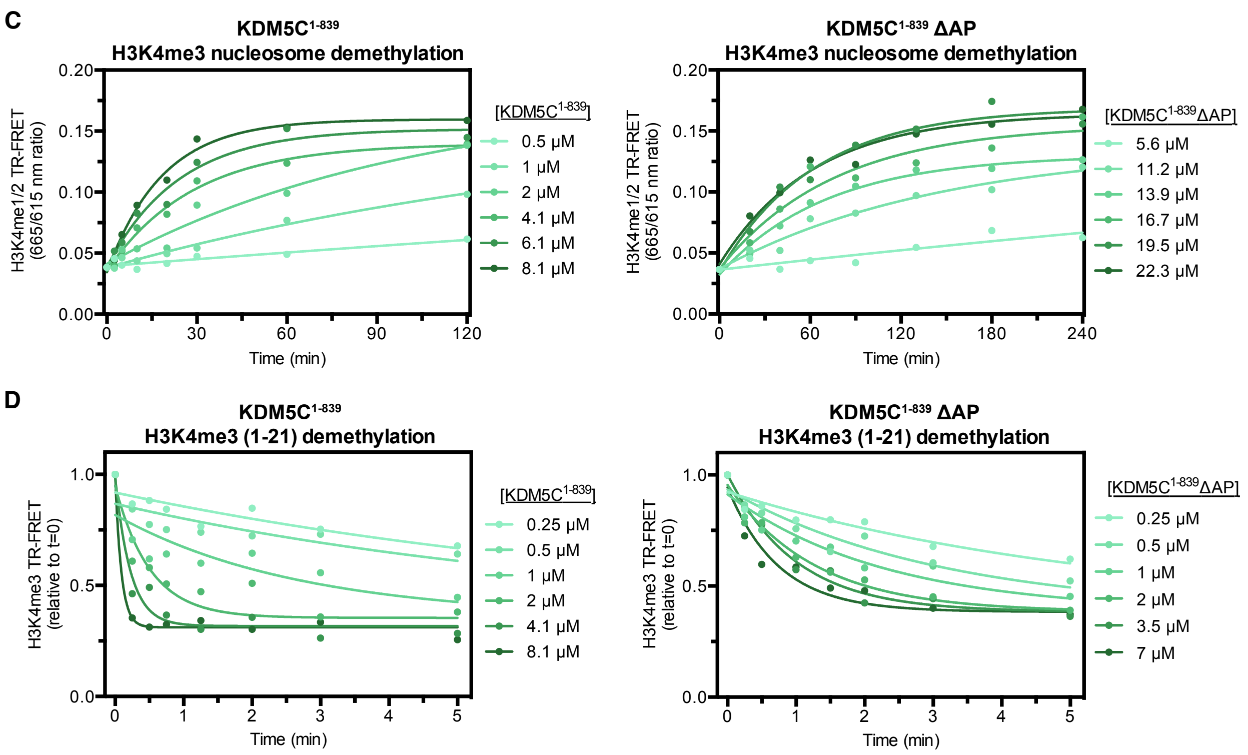


**Figure S1. Related to Figure 1.**

**(A)** H3K4me3 substrate peptide demethylation by KDM5C constructs. *Left*: Demethylation kinetics of the H3K4me3 (1-21) substrate peptide by KDM5C constructs under single turnover conditions measured by a TR-FRET based kinetic assay. Observed rates are fit to a cooperative kinetic model, with *n* denoting the Hill coefficient. Representative kinetic traces used to determine observed demethylation rates are in Figure S1D. *Right*: Demethylation kinetics of the H3K4me3 (1-21) substrate peptide by KDM5C constructs under multiple turnover conditions measured by a formaldehyde release based kinetic assay. Deletion of the ARID and PHD1 region results in higher demethylase activity on the substrate peptide under multiple turnover conditions due to loss of substrate inhibition caused by this region. **(B)** Unmodified and substrate core nucleosome binding by KDM5C$^{1\text{-}839}$ and KDM5C$^{1\text{-}839}$ ΔAP. Nucleosome binding curves were measured by EMSA and fit to a cooperative binding model to determine apparent dissociation constants ($K_d^{app}$), with *n* denoting the Hill coefficient (*top*). Representative gel shifts of KDM5C binding to nucleosomes (*bottom*). Due to unattainable saturation of binding, a lower limit for the dissociation constant is presented for the unmodified nucleosome. **(C)** Representative demethylation kinetic traces of substrate nucleosome demethylation by KDM5C constructs (*left*: KDM5C$^{1\text{-}839}$, *right*: KDM5C$^{1\text{-}839}$ ΔAP) under single turnover conditions using TR-FRET based kinetic assay detecting formation of the H3K4me1/2 product nucleosome over time. Observed rates ($k_{obs}$) are obtained by fitting kinetic traces to an exponential function. **(D)** Representative demethylation kinetic traces of substrate peptide demethylation by KDM5C constructs (*left*: KDM5C$^{1\text{-}839}$, *right*: KDM5C$^{1\text{-}839}$ ΔAP) under single turnover conditions using TR-FRET based kinetic assay detecting loss of the H3K4me3 substrate peptide over time. Observed rates ($k_{obs}$) are obtained by fitting kinetic traces to an exponential function. All error bars represent SEM of at least two independent experiments ($n \geq 2$).

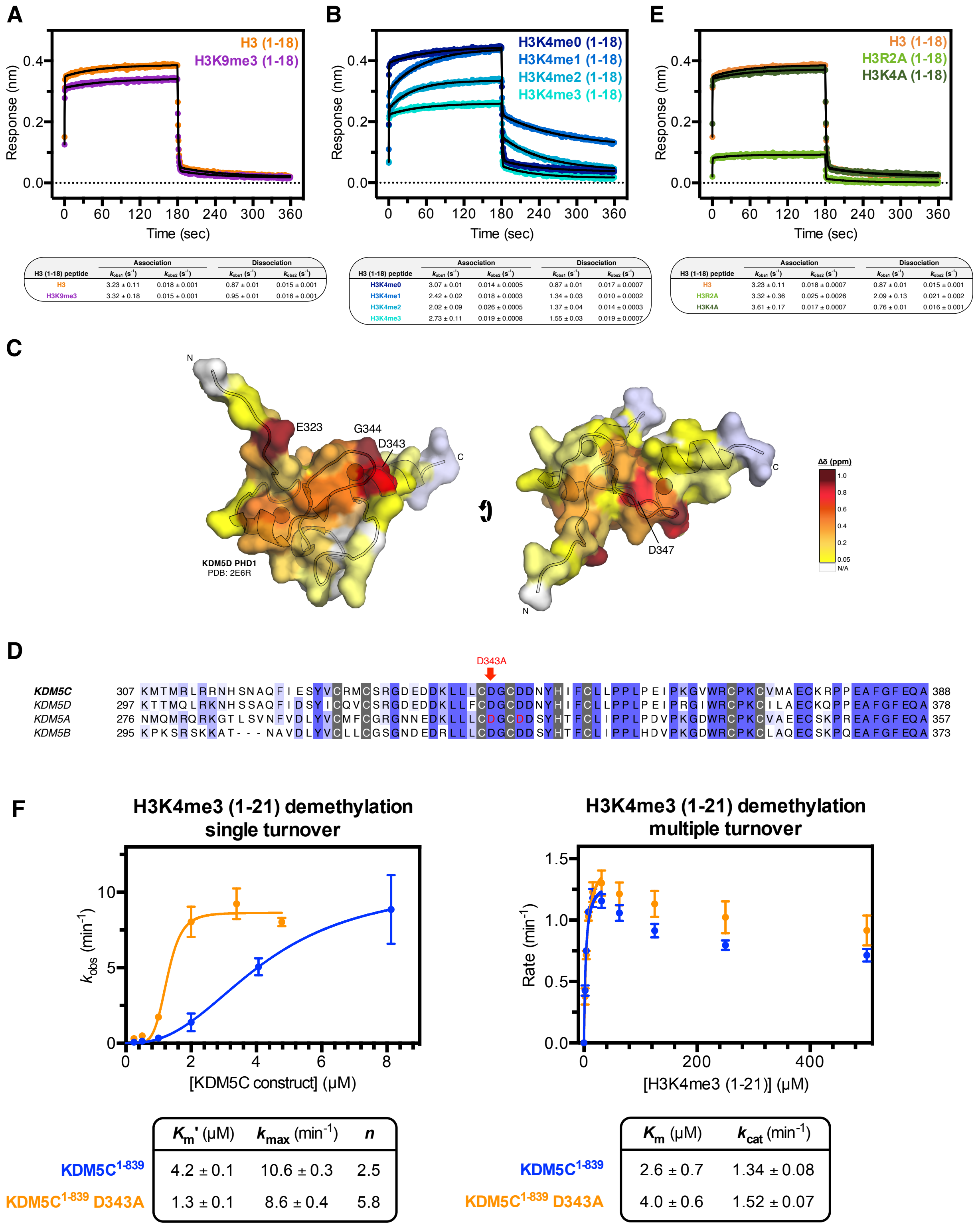


| H3 (1-18) peptide | Association | | Dissociation | |
|---|---|---|---|---|
| | $k_{obs1}$ ($s^{-1}$) | $k_{obs2}$ ($s^{-1}$) | $k_{obs1}$ ($s^{-1}$) | $k_{obs2}$ ($s^{-1}$) |
| H3 | 3.23 ± 0.11 | 0.018 ± 0.001 | 0.87 ± 0.01 | 0.015 ± 0.001 |
| H3K9me3 | 3.32 ± 0.18 | 0.015 ± 0.001 | 0.95 ± 0.01 | 0.016 ± 0.001 |

| H3 (1-18) peptide | Association | | Dissociation | |
|---|---|---|---|---|
| | $k_{obs1}$ ($s^{-1}$) | $k_{obs2}$ ($s^{-1}$) | $k_{obs1}$ ($s^{-1}$) | $k_{obs2}$ ($s^{-1}$) |
| H3K4me0 | 3.07 ± 0.01 | 0.014 ± 0.0005 | 0.87 ± 0.01 | 0.017 ± 0.0007 |
| H3K4me1 | 2.42 ± 0.02 | 0.018 ± 0.0003 | 1.34 ± 0.03 | 0.010 ± 0.0002 |
| H3K4me2 | 2.02 ± 0.09 | 0.026 ± 0.0005 | 1.37 ± 0.04 | 0.014 ± 0.0003 |
| H3K4me3 | 2.73 ± 0.11 | 0.019 ± 0.0008 | 1.55 ± 0.03 | 0.019 ± 0.0007 |

| H3 (1-18) peptide | Association | | Dissociation | |
|---|---|---|---|---|
| | $k_{obs1}$ ($s^{-1}$) | $k_{obs2}$ ($s^{-1}$) | $k_{obs1}$ ($s^{-1}$) | $k_{obs2}$ ($s^{-1}$) |
| H3 | 3.23 ± 0.11 | 0.018 ± 0.0007 | 0.87 ± 0.01 | 0.015 ± 0.001 |
| H3R2A | 3.32 ± 0.36 | 0.025 ± 0.0026 | 2.09 ± 0.13 | 0.021 ± 0.002 |
| H3K4A | 3.61 ± 0.17 | 0.017 ± 0.0007 | 0.76 ± 0.01 | 0.016 ± 0.001 |

| | $K_m$' (μM) | $k_{max}$ (min$^{-1}$) | $n$ |
|---|---|---|---|
| KDM5C$^{1\text{-}839}$ | 4.2 ± 0.1 | 10.6 ± 0.3 | 2.5 |
| KDM5C$^{1\text{-}839}$ D343A | 1.3 ± 0.1 | 8.6 ± 0.4 | 5.8 |

| | $K_m$ (μM) | $k_{cat}$ (min$^{-1}$) |
|---|---|---|
| KDM5C$^{1\text{-}839}$ | 2.6 ± 0.7 | 1.34 ± 0.08 |
| KDM5C$^{1\text{-}839}$ D343A | 4.0 ± 0.6 | 1.52 ± 0.07 |

**G**

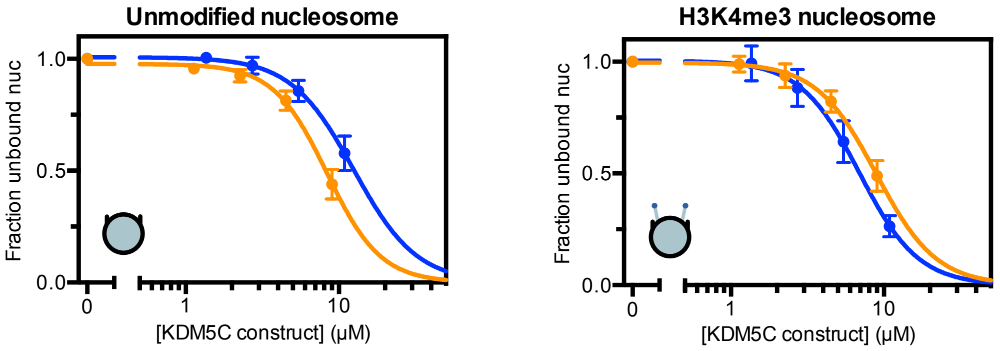


| | $K_d$ app | $n$ |
|---|---|---|
| KDM5C[1-839] | ≥ 12.5 ± 1.2 μM | 2.1 |
| KDM5C[1-839] D343A | 8.4 ± 0.5 μM | 2.5 |

| | $K_d$ app | $n$ |
|---|---|---|
| KDM5C[1-839] | 6.9 ± 0.8 μM | 2.3 |
| KDM5C[1-839] D343A | 8.9 ± 0.7 μM | 2.2 |

**H**

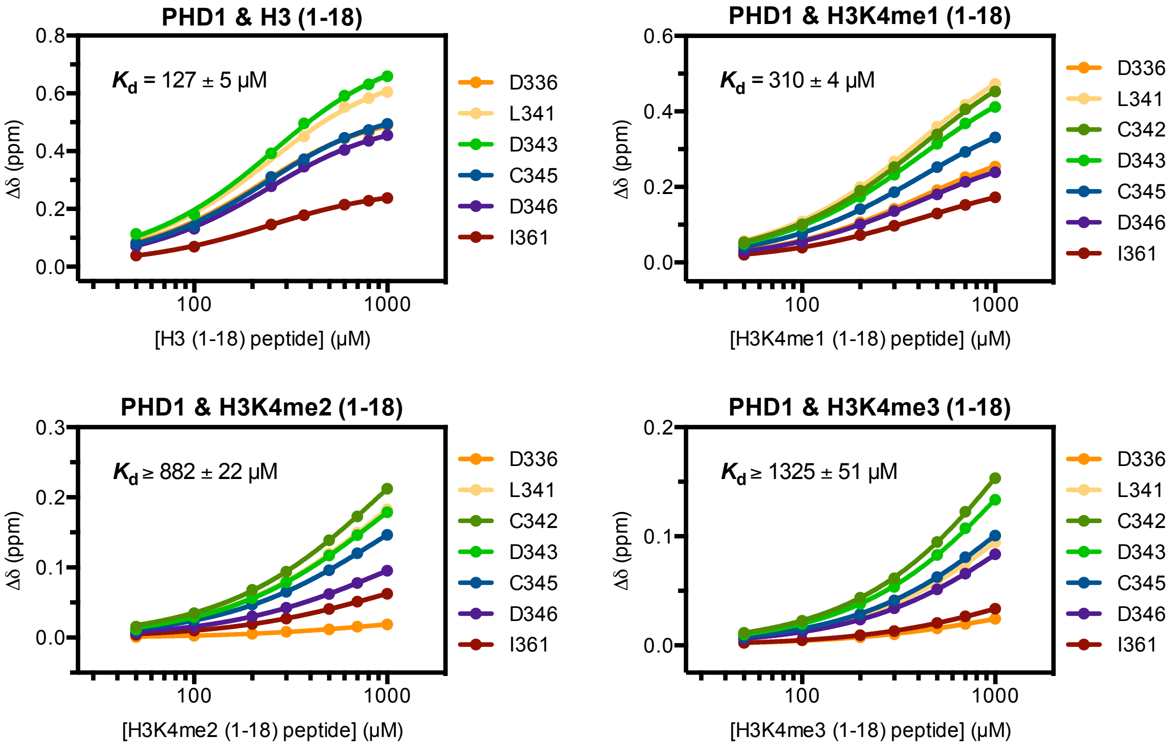


**Figure S2. Related to Figure 2.**
**(A)** Binding kinetic trace of immobilized Avitag-PHD1 binding to H3 (1-18) and H3K9me3 (1-18) tail peptides measured by bio-layer interferometry (BLI). Observed rates ($k_{obs}$) of association and dissociation are obtained by fitting kinetic traces to a two phase exponential function. **(B)** Binding kinetic trace of immobilized Avitag-PHD1 binding to H3K4me0/1/2/3 (1-18) tail peptides measured by BLI. Biphasic kinetic binding by PHD1 is modulated by the H3K4me state. **(C)** Chemical shift perturbations of PHD1 residues upon binding of the H3 (1-18) tail peptide (Figure 2A) colored by the gradient, unperturbed (yellow) to significantly perturbed (maroon), mapped to homologous residues in KDM5D PHD1 structure (PDB: 2E6R). Significantly perturbed residues are labeled. **(D)** Binding kinetic trace of immobilized Avitag-PHD1 binding to H3 (1-18) and H3 mutant (1-18) tail peptides (H3R2A and H3K4A) measured by BLI. Recognition of the H3 tail by PHD1 depends on the R2 residue but not K4 residue in H3. **(E)** Sequence alignment of PHD1 domains in KDM5A-D. The H3R2 recognizing residues D312 and D315 of KDM5A are indicated in red, and the PHD1 mutation D343A from this study is denoted above KDM5C. Zinc coordinating residues

are highlighted in gray. **(F)** H3K4me3 substrate peptide demethylation by PHD1 mutant KDM5C$^{1-839}$ relative to wild type. *Left*: Demethylation kinetics of the H3K4me3 (1-21) substrate peptide under single turnover conditions measured by a TR-FRET based kinetic assay. Observed rates are fit to a cooperative kinetic model, with *n* denoting the Hill coefficient. Unlike on the substrate nucleosome, the D343A PHD1 mutation does not increase catalytic rate on the substrate peptide, but does increase overall catalytic efficiency. *Right*: Demethylation kinetics of the H3K4me3 (1-21) substrate peptide under multiple turnover conditions measured by a formaldehyde release based kinetic assay. The D343A PHD1 mutation does not affect catalysis on the substrate peptide under these conditions, nor does it significantly affect substrate inhibition. **(G)** Unmodified and substrate core nucleosome binding by PHD1 mutant KDM5C$^{1-839}$ relative to wild type. Nucleosome binding curves were measured by EMSA and fit to a cooperative binding model to determine apparent dissociation constants ($K_d^{app}$), with *n* denoting the Hill coefficient. Due to unattainable saturation of binding, a lower limit for the dissociation constant is presented for WT KDM5C binding the unmodified nucleosome. **(H)** Binding of the H3K4me0/1/2/3 (1-18) tail peptides by PHD1 by NMR titration HSQC experiments of indicated PHD1 residues that localize to the H3 binding surface (Figure S2C). Average dissociation constants with standard error for each ligand were determined from dissociation constants obtained from chemical shift changes (Δδ) of individual PHD1 residues. Due to incomplete saturation of binding, a lower limit for the dissociation constant is presented for the H3K4me2/3 peptides. All error bars represent SEM of at least two independent experiments ($n \geq 2$).

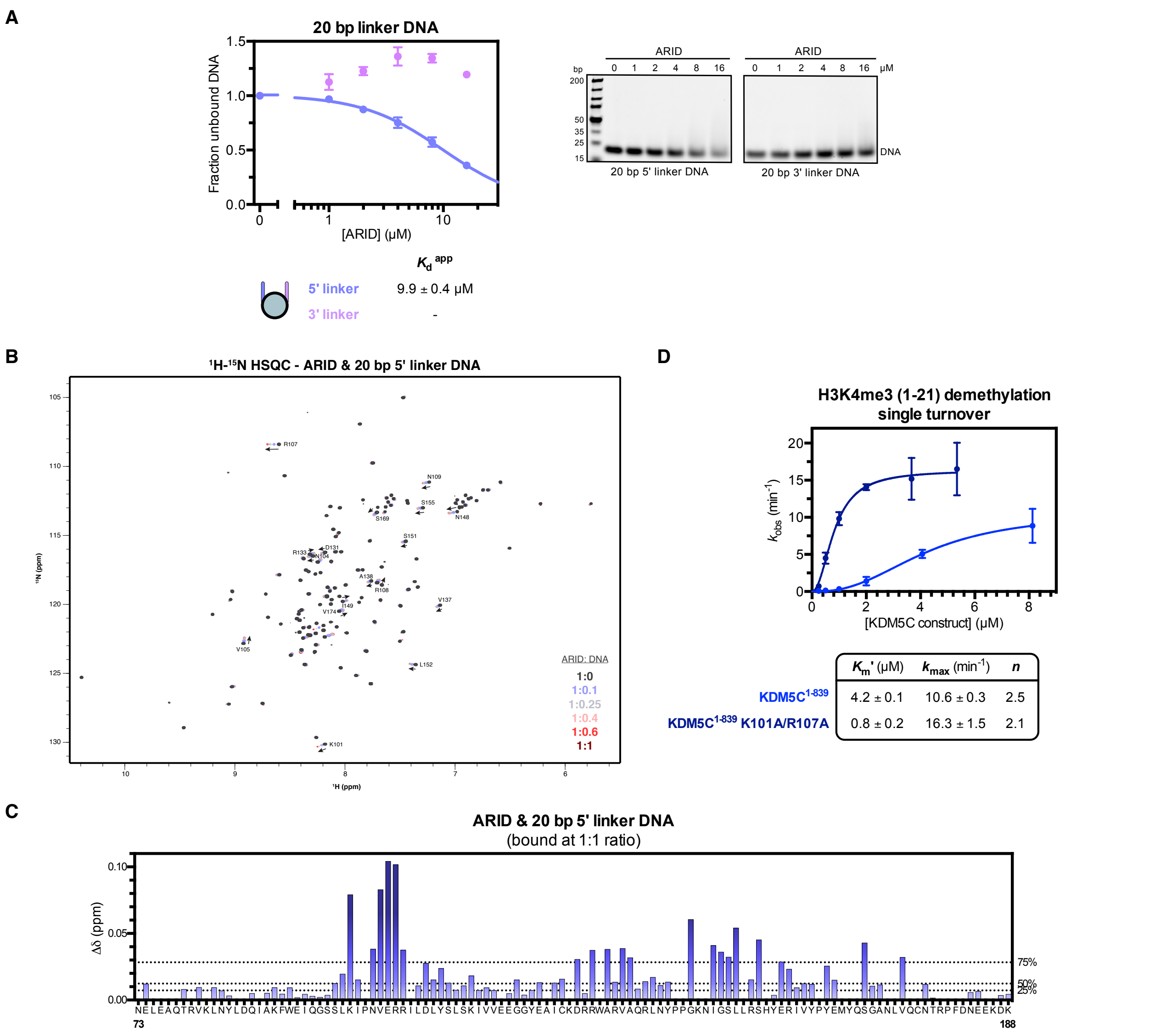


**Figure S3. Related to Figure 3.**

**(A)** 20 bp linker DNA fragment binding by the ARID domain. Fragments contain 5' and 3' flanking DNA sequences used in the 187 bp nucleosome. Binding curves were measured by EMSA and fit to a binding model to determine apparent dissociation constants ($K_d^{app}$) (*left*). Representative gel shifts of ARID binding to 20 bp flanking linker DNA fragments (*right*). **(B)** 2D $^1$H-$^{15}$N HSQC spectra of ARID titrated with increasing amounts of the 5' linker DNA 20 bp fragment with indicated molar ratios. Assignments of most perturbed residues in ARID are labeled. **(C)** Chemical shift change (Δδ) of ARID residue backbone assignments upon binding of the 5' linker DNA 20 bp fragment at 1:1 molar ratio measured by NMR. ARID backbone assignments could not be reliably transferred to a subset of residues and thus chemical shifts could not be determined (indicated by no values). Dashed lines indicate 25th, 50th, and 75th percentile rankings, and residues are colored by a gradient from unperturbed (light blue) to significantly perturbed (navy). **(D)** Demethylation kinetics of the H3K4me3 (1-21) substrate peptide by wild type and ARID mutant KDM5C$^{1-839}$ under single turnover conditions. Observed rates are fit to a cooperative kinetic model, with $n$ denoting the Hill coefficient. Unlike on the substrate nucleosome, the K101A/R107A ARID double mutation does not decrease catalytic rate on the substrate peptide, but does increase overall catalytic efficiency. All error bars represent SEM of at least two independent experiments ($n \geq 2$).

**A**

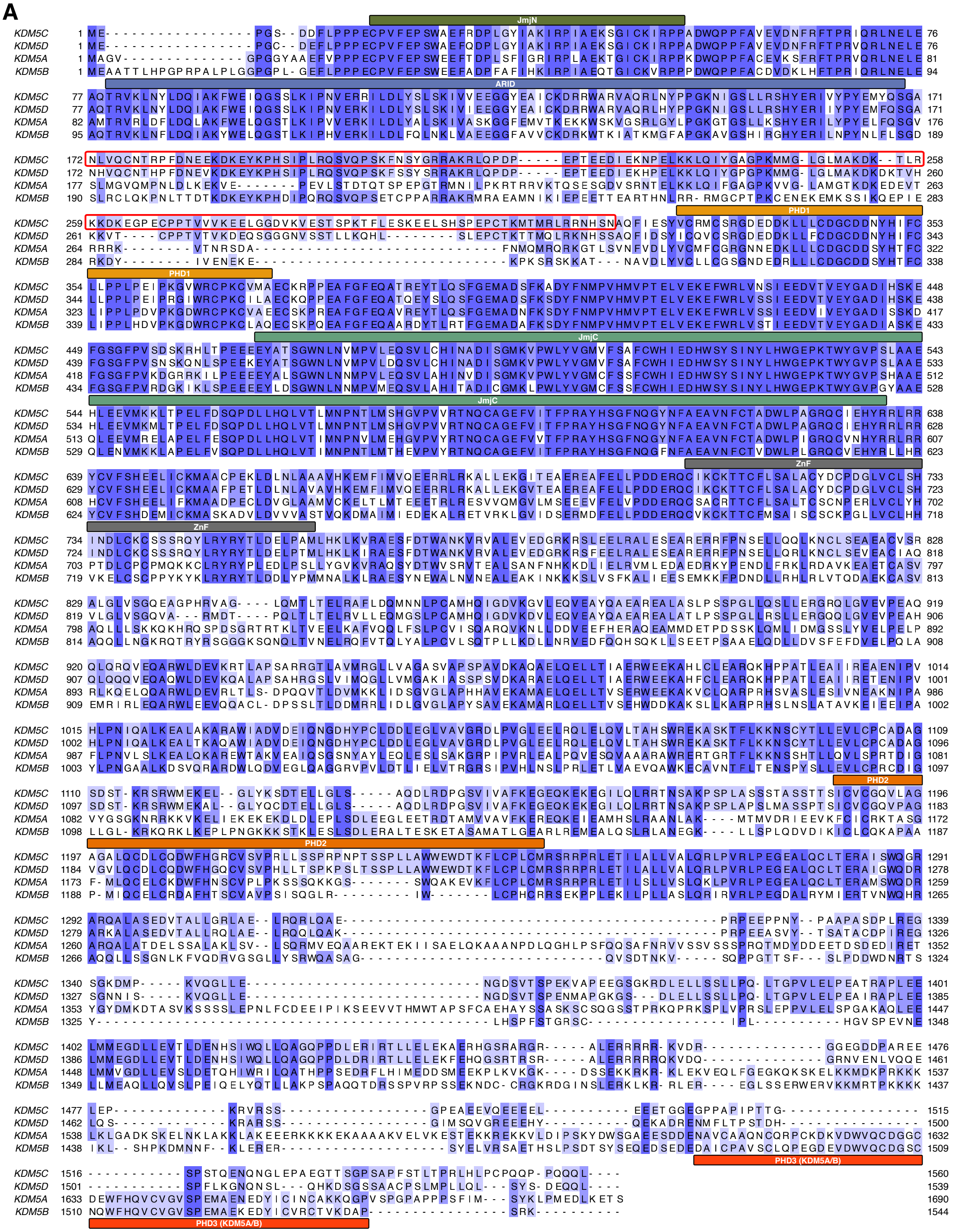
JmjN
ARID
PHD1
JmjC
ZnF
PHD2
PHD3 (KDM5A/B)
KDM5C
KDM5D
KDM5A
KDM5B

**B**

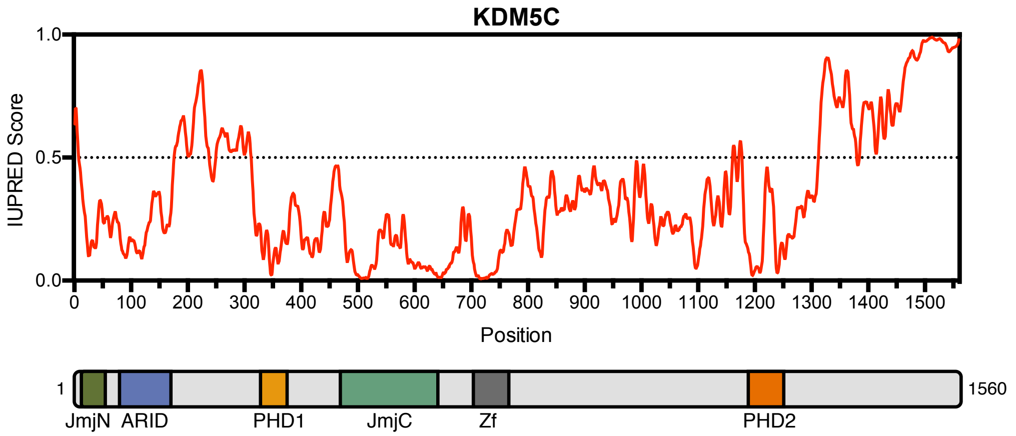


**C**

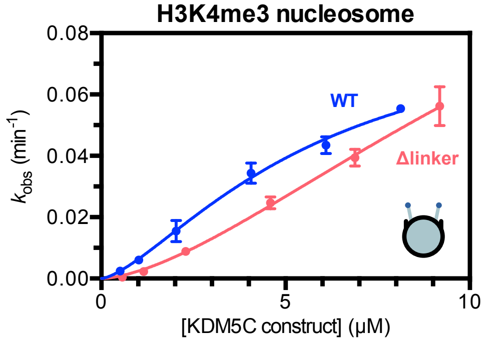


| | $K_m^{app}$ (μM) | $k_{max}$ (min$^{-1}$) | $n$ |
|---|---|---|---|
| KDM5C[1-839] | 5.7 ± 1.7 | 0.087 ± 0.018 | 1.5 |
| KDM5C[1-839] Δlinker | 14.2 ± 5.3 | 0.168 ± 0.061 | 1.6 |

**D**

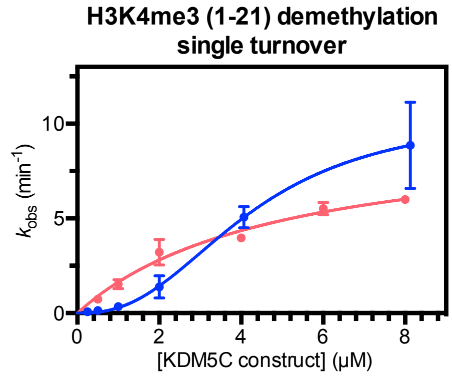


| | $K_m'$ (μM) | $k_{max}$ (min$^{-1}$) | $n$ |
|---|---|---|---|
| KDM5C[1-839] | 4.2 ± 0.1 | 10.6 ± 0.3 | 2.5 |
| KDM5C[1-839] Δlinker | 4.9 ± 1.3 | 9.7 ± 1.3 | 1.0 |

**E**

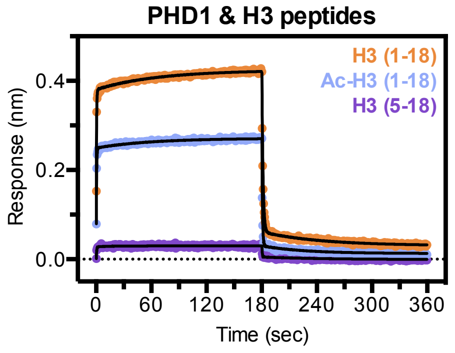


| | Association | | Dissociation | |
|---|---|---|---|---|
| H3 peptide | $k_{obs1}$ (s$^{-1}$) | $k_{obs2}$ (s$^{-1}$) | $k_{obs1}$ (s$^{-1}$) | $k_{obs2}$ (s$^{-1}$) |
| H3 (1-18) | 2.79 ± 0.08 | 0.015 ± 0.0006 | 0.84 ± 0.01 | 0.015 ± 0.0009 |
| Ac-H3 (1-18) | 2.51 ± 0.08 | 0.016 ± 0.0011 | 1.63 ± 0.03 | 0.022 ± 0.0016 |
| H3 (5-18) | 2.14 ± 0.26 | 0.063 ± 0.0217 | 2.34 ± 0.48 | 0.015 ± 0.0040 |

**H**

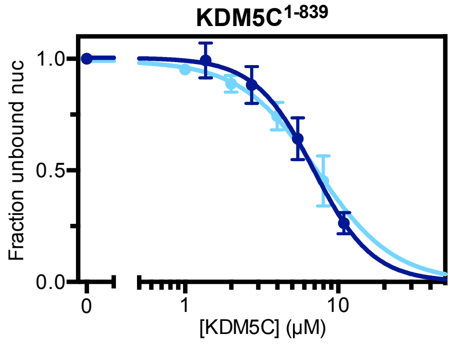


| | $K_d^{app}$ | $n$ |
|---|---|---|
| 147 bp H3K4me3 nuc | 6.9 ± 0.8 μM | 2.3 |
| 187 bp H3K4me3 nuc | 7.3 ± 0.9 μM | 1.7 |

**F**

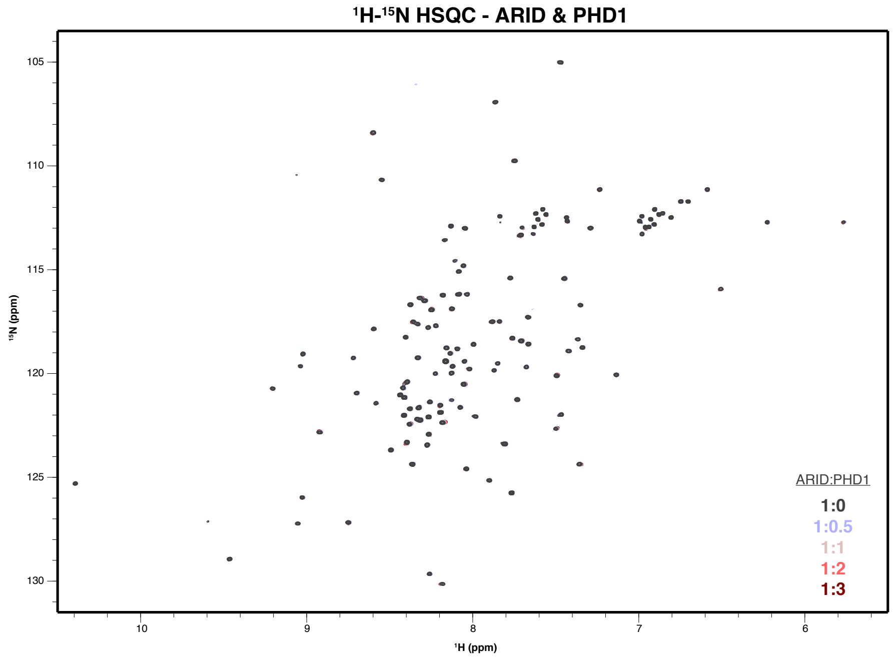

¹H-¹⁵N HSQC - ARID & PHD1
¹⁵N (ppm)
¹H (ppm)
ARID:PHD1
1:0
1:0.5
1:1
1:2
1:3


**G**

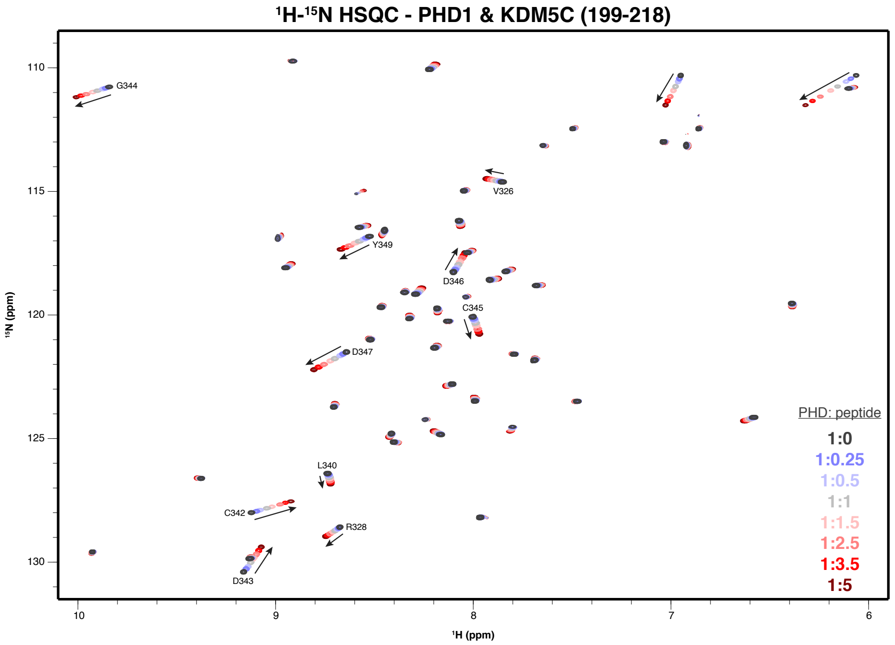

¹H-¹⁵N HSQC - PHD1 & KDM5C (199-218)
G344
V326
Y349
D346
C345
D347
L340
C342
R328
D343
¹⁵N (ppm)
¹H (ppm)
PHD: peptide
1:0
1:0.25
1:0.5
1:1
1:1.5
1:2.5
1:3.5
1:5


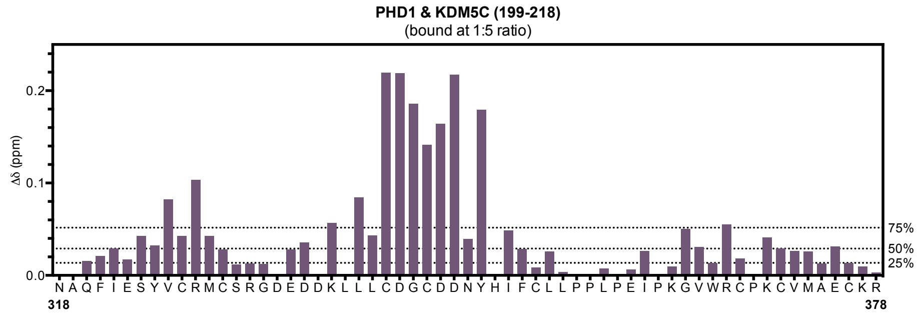

PHD1 & KDM5C (199-218)
(bound at 1:5 ratio)
Δδ (ppm)
75%
50%
25%
N A Q F I E S Y V C R M C S R G D E D D K L L L C D G C D D N Y H I F C L L P P L P E I P K G V W R C P K C V M A E C K R
318
378

**I**

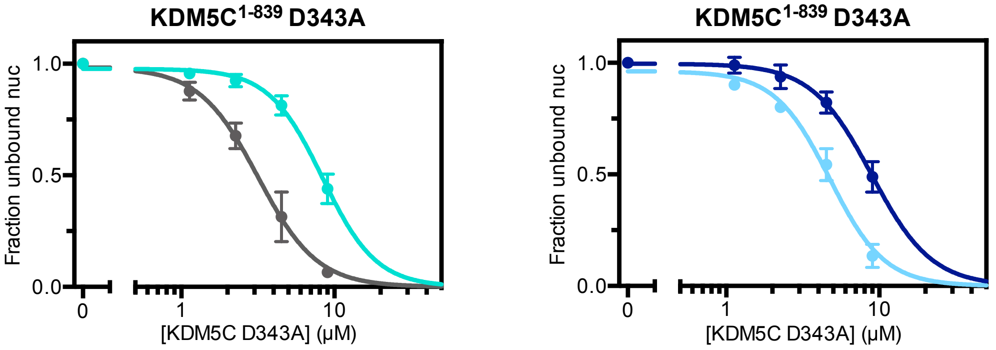


| | $K_d^{app}$ | $n$ | | $K_d^{app}$ | $n$ |
|---|---|---|---|---|---|
| 147 bp nuc | 8.4 ± 0.5 μM | 2.5 | 147 bp H3K4me3 nuc | 8.9 ± 0.7 μM | 2.2 |
| 187 bp nuc | 3.2 ± 0.4 μM | 2.3 | 187 bp H3K4me3 nuc | 4.7 ± 0.3 μM | 2.4 |

**Figure S4. Related to Figure 4.**
**(A)** Sequence alignment of human KDM5A-D with annotated domains. KDM5C has a different and extended linker region between ARID and PHD1 (boxed in red). **(B)** IUPred profile [74] of predicted disorder in KDM5C (*top*) and annotated domain architecture of KDM5C (*bottom*). The linker between ARID and PHD1 is predicted to be disordered. **(C)** Demethylation kinetics of the H3K4me3 substrate nucleosome by wild type and $KDM5C^{1-839}$ Δlinker under single turnover conditions. Observed rates are fit to a cooperative kinetic model, with $n$ denoting the Hill coefficient. Deletion of the ARID-PHD1 linker does not significantly affect the catalytic efficiency of substrate nucleosome demethylation. **(D)** Demethylation kinetics of the H3K4me3 (1-21) substrate peptide by wild type and $KDM5C^{1-839}$ Δlinker under single turnover conditions. Observed rates are fit to a cooperative kinetic model, with $n$ denoting the Hill coefficient. Similarly to nucleosomes, deletion of the ARID-PHD1 linker does not significantly affect the catalytic efficiency of substrate peptide demethylation. **(E)** Binding kinetic trace of immobilized Avitag-PHD1 binding to H3 (1-18), N-terminally acetylated H3 (1-18), and H3 (5-18) tail peptides measured by bio-layer interferometry. Observed rates ($k_{obs}$) of association and dissociation are obtained by fitting kinetic traces to a two phase exponential function. Recognition of the H3 tail by PHD1 does not strongly depend on the H3 N-terminus but does depend on the first 4 residues of H3 (ARTK). **(F)** 2D $^1H$-$^{15}N$ HSQC spectra of ARID titrated with increasing amounts of PHD1 with indicated molar ratios. PHD1 does not appear to bind the ARID domain. **(G)** 2D $^1H$-$^{15}N$ HSQC spectra of PHD1 titrated with increasing amounts of KDM5C (199-218) peptide with indicated molar ratios (*top*). Assignments of most perturbed residues in PHD1 are labeled. Corresponding chemical shift change (Δδ) of PHD1 residues upon binding of the KDM5C (199-218) peptide at 1:5 molar ratio (PHD:peptide) (*bottom*). Dashed lines indicate 25th, 50th, and 75th percentile rankings. **(H)** Nucleosome binding curves of $KDM5C^{1-839}$ binding to substrate nucleosomes with and without 20 bp flanking DNA. Nucleosome binding curves were measured by EMSA and fit to a cooperative binding model to determine apparent dissociation constants ($K_d^{app}$), with $n$ denoting the Hill coefficient. **(I)** Binding curves of PHD1 mutant $KDM5C^{1-839}$ binding to unmodified and substrate nucleosomes with and without 20 bp flanking DNA. All error bars represent SEM of at least two independent experiments ($n \geq 2$).

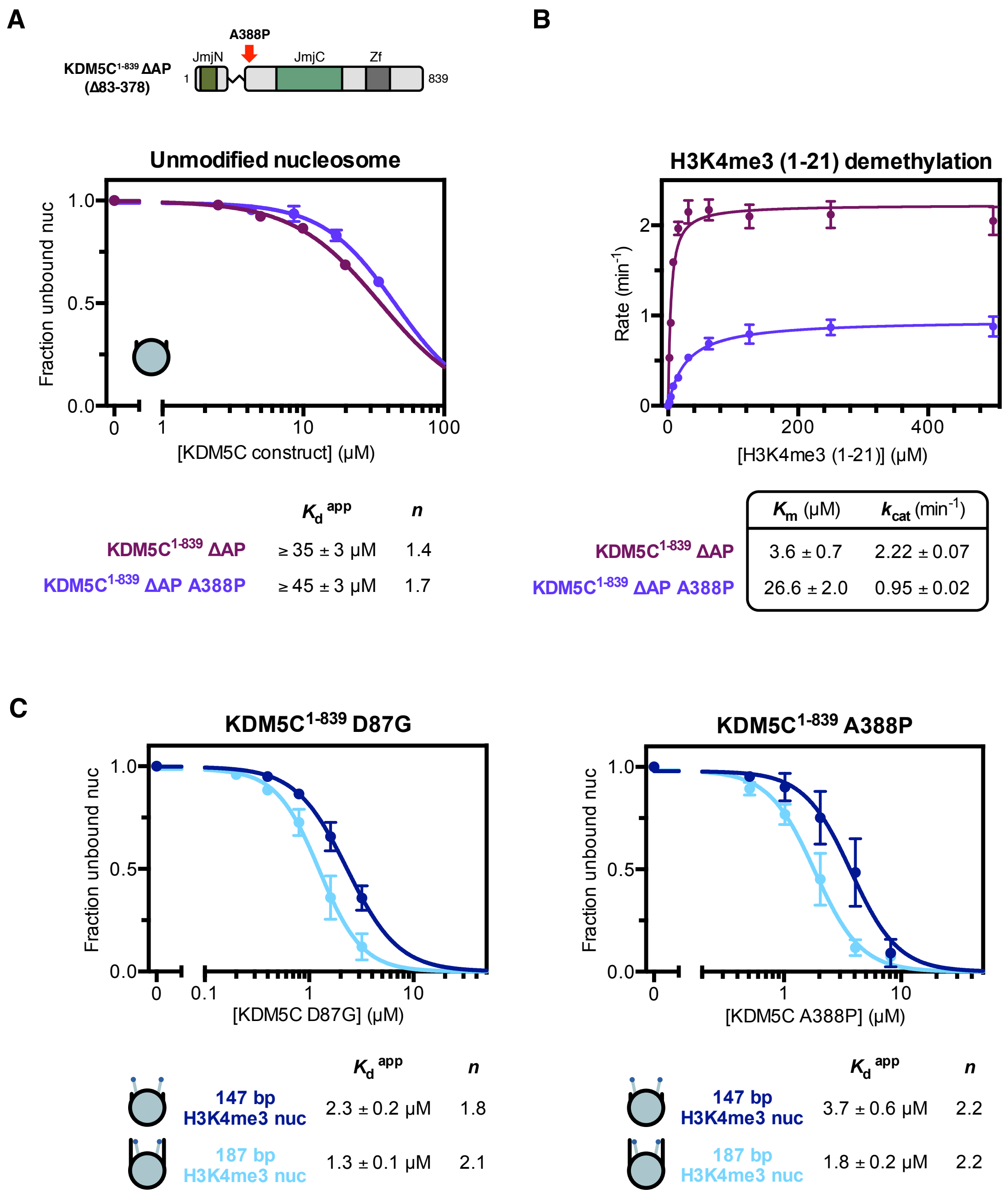


**Figure S5. Related to Figure 5.**

**(A)** Unmodified core nucleosome binding by KDM5C$^{1\text{-}839}$ ΔAP wild type and A388P. Nucleosome binding curves were measured by EMSA and fit to a cooperative binding model to determine apparent dissociation constants ($K_d^{app}$), with $n$ denoting the Hill coefficient. Due to unattainable saturation of binding, a lower limit for the dissociation constant is presented. The A388P mutation does not enhance nucleosome binding in the absence of the ARID and PHD1 region, indicating this region in KDM5C is altered by the A388P mutation to enable enhanced binding. **(B)** Demethylation kinetics of the H3K4me3 (1-21) substrate peptide by KDM5C$^{1\text{-}839}$ ΔAP wild type and A388P under multiple turnover conditions measured by a formaldehyde release based kinetic assay. The A388P mutation reduces demethylase activity of the catalytic domain alone, indicating distal structural disruption of the catalytic domain by this mutation. **(C)** Binding curves of KDM5C$^{1\text{-}839}$ D87G and A388P binding to substrate nucleosomes with and without 20 bp flanking DNA. All error bars represent SEM of at least two independent experiments ($n \geq 2$).
